# High-resolution structural analysis of enterovirus-reactive polyclonal antibodies in complex with whole virions

**DOI:** 10.1101/2022.01.31.478566

**Authors:** Aleksandar Antanasijevic, Autumn J. Schulze, Vijay S. Reddy, Andrew B. Ward

## Abstract

Non-polio enteroviruses (NPEVs) cause serious illnesses in young children and neonates including aseptic meningitis, encephalitis, neonatal sepsis and inflammatory muscle disease, among others. Over 100 serotypes have been described to date but except for the EV-A71, there are no available vaccines or therapeutics against NPEVs. Efforts towards rationally designed pan-NPEV vaccines would greatly benefit from structural biology methods for rapid and comprehensive evaluation of vaccine candidates and elicited antibody responses. Towards this goal, we tested if electron-microscopy-based polyclonal epitope mapping (EMPEM), a method where structural analysis is performed using serum-derived polyclonal antibodies (pAbs), can be applied to an NPEV. EMPEM was performed on immune complexes featuring CV-A21 viral particles and CV-A21-specific pAbs isolated from mice. Notably, this is the first example of structural analysis of polyclonal immune complexes comprising whole virions. We introduce a data processing workflow that allows reconstruction of different pAbs at near-atomic resolution. The analysis resulted in identification of several antibodies targeting two immunodominant epitopes, near the 3-fold and 5-fold axis of symmetry; the latter overlapping with the receptor binding site. These results demonstrate that EMPEM can be applied to map broad-spectrum polyclonal immune responses against intact virions and define potentially cross-reactive epitopes.

## Introduction

Non-polio enteroviruses (NPEVs) from the family of *Picornaviridae* are non-enveloped viruses composed of (+) ssRNA genomes that infect humans and cause a variety of human diseases (Granoff & Webster, 1999; Nikonov, Chernykh, Garber, & Nikonova, 2017). Rhinoviruses (RVs), coxsackieviruses (CVs), echoviruses and recently emergent EV-A71 and EV-D68 are examples of viruses belonging to this family. NPEVs account for 10-15 million infections each year, particularly among neonates and young children, that result in various illnesses ranging from aseptic meningitis, viral encephalitis, acute diarrhea with echovirus 30 (E30), hand, foot and mouth disease and severe respiratory illness to inflammatory muscle disease that is closely associated with acute flaccid myelitis (Murphy et al., 2021; Oyero, Adu, & Ayukekbong, 2014). There are currently over 100 different serotypes of NPEVs that have been identified and these numbers are likely to increase due to the high intrinsic mutation rates during the viral replication as well as greater recombination rates between different strains. Despite a relatively large number of circulating NPEVs, no effective vaccines or therapeutics are available except for the EV-A71 vaccine approved for use in some Asian countries (Puenpa, Wanlapakorn, Vongpunsawad, & Poovorawan, 2019). There is a need for new therapy and vaccine solutions, particularly the ones capable of targeting multiple NPEVs instead of each one individually.

Structural biology methods, such as cryo-electron microscopy (cryoEM), have been widely used for characterization of vaccine candidates and vaccine-elicited monoclonal antibodies (mAbs) (Li, Li, Fu, & Cao, 2020; Stass, Ilca, & Huiskonen, 2018; Ward & Wilson, 2017). However, these studies have been impeded by the relatively low throughput (one sample can be analyzed at a time) and the need for mAb isolation. EM-based polyclonal epitope mapping (EMPEM) is a tool for rapid and comprehensive structural analysis of immune complexes featuring vaccine-elicited polyclonal antibodies (pAbs) purified from sera (Bianchi et al., 2018)(Antanasijevic et al., 2021; Bangaru et al., 2021; Nogal et al., 2020). Additionally, we have recently shown that structural information from the reconstructed high-resolution maps can be used to identify sequences of polyclonal antibody families (Antanasijevic et al., 2022). EMPEM has thus far been only applied to recombinantly-produced antigens. In the case of enteroviruses this is not possible, given the difficulty of producing individual capsid components, nor is it desirable, because the majority of elicited antibodies have been found to target quaternary epitopes on the viral surface (He et al., 2020; Xu et al., 2017; Xu et al., 2021).

Herein, we introduce an EMPEM pipeline for rapid structural analysis of pAb responses elicited by enteroviruses using whole viral particles. Our data processing workflow enabled reconstruction of immune complexes featuring pAbs at near atomic resolution. In a case study using the Coxsackievirus A21 (CV-A21), we discovered two immunodominant sites on the surface of CV-A21 readily targeted by antibodies; one in immediate proximity to the receptor binding site. Overall, we demonstrate the feasibility of this approach and provide valuable insights for future vaccine design efforts.

## Results

For this proof-of-concept study we selected CV-A21 viral strain responsible for a substantial proportion of enterovirus-associated acute respiratory tract infections in humans (Couch, Douglas, Lindgren, Gerone, & Knight, 1970; Zou et al., 2017). Additionally, this virus is evaluated clinically for its oncolytic potential (Bradley et al., 2014; McCarthy, Jayawardena, Burga, & Bostina, 2019). CV-A21 and other NPEVs share the same overall icosahedral capsid structure consisting of 60 copies of VP1-4 that are arranged with pseudo, T*=3* symmetry (Figure 1A) (Basavappa et al., 1994; Curry et al., 1997; Xiao et al., 2005). Each of the VP1-3 protein subunits displays compact tertiary structure comprising a jelly-roll b-sandwich, which form the bulk of the capsid and the outer surface, making these subunits the primary targets for antibodies (Figure 1A). VP4 adopts an extended conformation and is largely located on the inner surface of the capsid.

**Figure 1.**
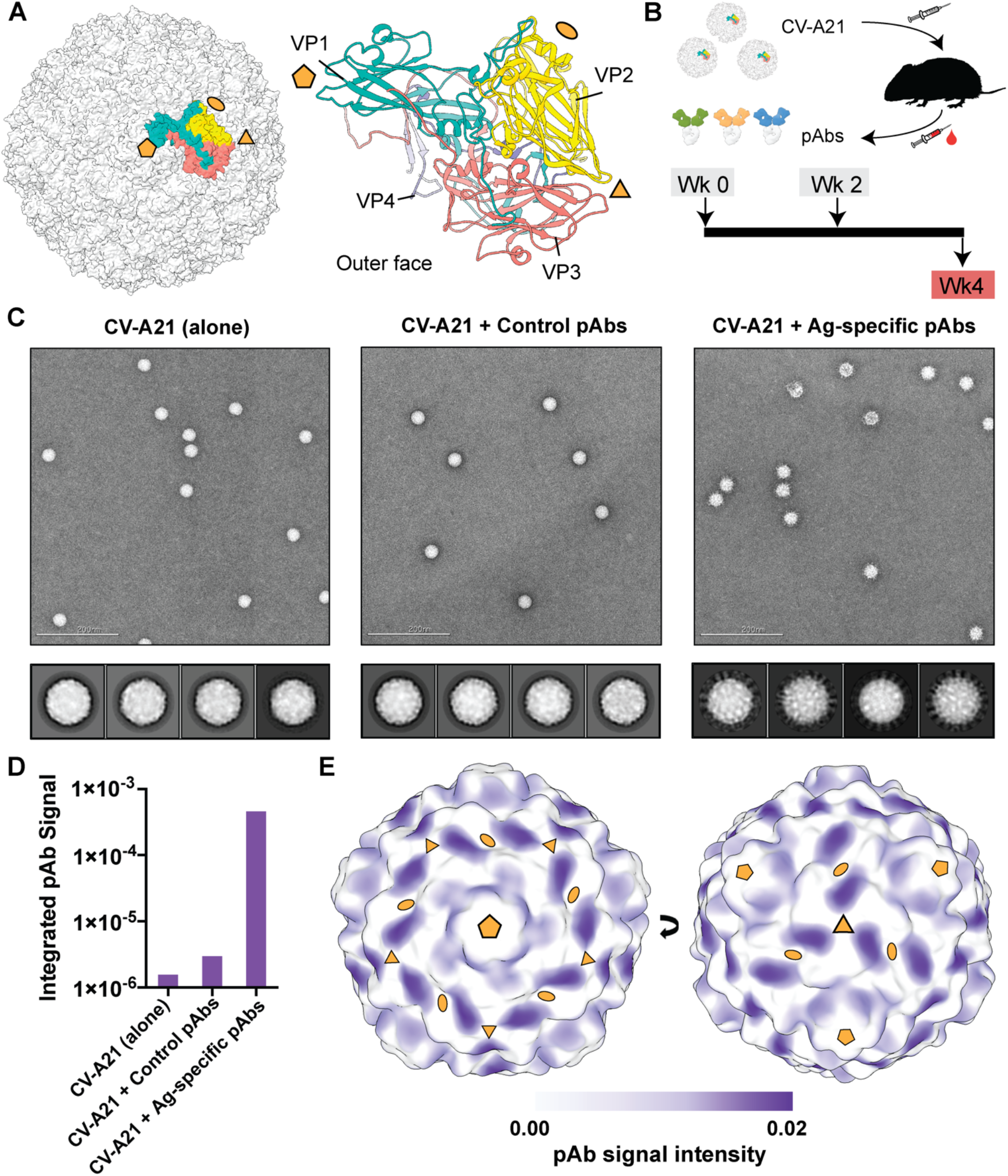
Immunization experiments and nsEMPEM analysis of polyclonal immune complexes. **[A]** Assembly of the CV capsid (left) and structure of the asymmetric unit (right). CV-A21 structure (PDB ID 1z7s; (Xiao et al., 2005)) used to make the figure. VP1-4 are colored differently and labeled for clarity. **[B]** Schematic representation of the workflow used to generate pAb samples (top) and immunization regimen details (bottom). **[C]** Representative EM micrograph (top) and 2D class averages (bottom) of CV-A21 viral particles alone (left) or in complex with pAbs (as Fab fragments) isolated from CV-A21-naïve mice (middle) or mice immunized with CV-A21 (right). **[D]** Integrated pAb-corresponding signal in 3D maps reconstructed using the nsEM data presented in panel C. This data is generated using Volume Tools in UCSF Chimera (Pettersen et al., 2004). **[E]** pAb-corresponding signal plotted on the surface of low-pass filtered map of CV-A21 viral capsid. Higher signal intensity corresponds to higher pAb occupancy. The icosahedral symmetry axes in panels A and E are indicated with golden-yellow pentamers (5-fold), triangles (3-fold) and ovals (2-fold).

We sought to structurally characterize the immunogenicity of CV-A21 virus and identify primary immunogenic epitopes using a mouse animal model. Purified CV-A21 viral particles (Figure 1 – figure supplement 1A,B) were injected into mice (Figure 1B). A control group of mice received PBS vehicle at the corresponding time points. Serum was harvested at week 4, following two antigen doses. For structural characterization, we isolated polyclonal antibodies from pooled, heat-inactivated mouse serum and cleaved them into Fab and Fc fragments with papain. Digested Fab/Fc samples were complexed with formaldehyde-treated CV-A21 particles and subjected to a round of size exclusion chromatography (SEC) to purify the viral fraction from unbound Fab/Fc (Figure 1 – figure supplement 1C). Inactivation with formaldehyde was necessary to execute experiments under biosafety level 1 conditions.

Assembled immune complexes (or free virions) were first characterized using negative stain EM (nsEMPEM; Figure 1C). pAbs isolated from the control group of mice did not bind to CV-A21, based on the combined analysis of the SEC data, raw EM images and 2D class averages. Conversely, antibodies isolated from animals infected with CV-A21 bound the antigen, as evidenced by the strong shift of virus-corresponding peak in SEC (Figure 1 – figure supplement 1C) and a “cloud” of pAbs (as Fab fragments) surrounding viral particles in EM images and 2D class averages (Figure 1C, right).

We reconstructed 3D maps from particles in the abovementioned datasets and performed partial signal integration in UCSF Chimera (Figure 1D, Figure 1 – figure supplement 1D; (Pettersen et al., 2004)). pAb-corresponding signal in the polyclonal sample from CV-A21-infected mice was higher by ~2 orders of magnitude compared to the control pAb sample, further supporting the observations made on the level of raw images and 2D. 3D maps were further used to map epitopes on CV-A21 particle targeted by pAbs (Figure 1E, Figure 1 – figure supplement 1D). Normalized map signal above noise threshold was observed at 2 sites, near the 3-fold symmetry axis (Site-1) and near the 5-fold symmetry axis (Site-2). pAb cloud at Site-2 is more diffuse and less intense compared to Site-1, indicating greater diversity in antibody response.

To further assess the diversity of polyclonal antibodies and precisely map epitope contacts, we subjected CV-A21 immune complexes to high-resolution cryoEM-based polyclonal epitope mapping (cryoEMPEM; (Antanasijevic et al., 2021)) (Figure 2A, Figure 2 - figure supplement 1 and 2). Briefly, the vitrified immune complexes were imaged using cryoEM and particles were aligned with icosahedral symmetry restraints imposed to obtain the initial reconstruction of the entire immune complex. Pre-aligned particles were then symmetry-expanded, thus appending all bound pAbs onto every protomer and increasing the initial dataset by 60-fold. This allowed positioning of solvent masks around different epitopes in a single protomer and classification of sub-particles based on structural properties of bound pAbs (i.e., to combine particles with structurally similar pAbs into unique subsets). Through iterative rounds of focused classification, we reconstructed maps featuring structurally unique polyclonal antibodies bound to epitopes at near atomic resolution (structurally unique polyclonal antibody class - pAbC).

**Figure 2.**
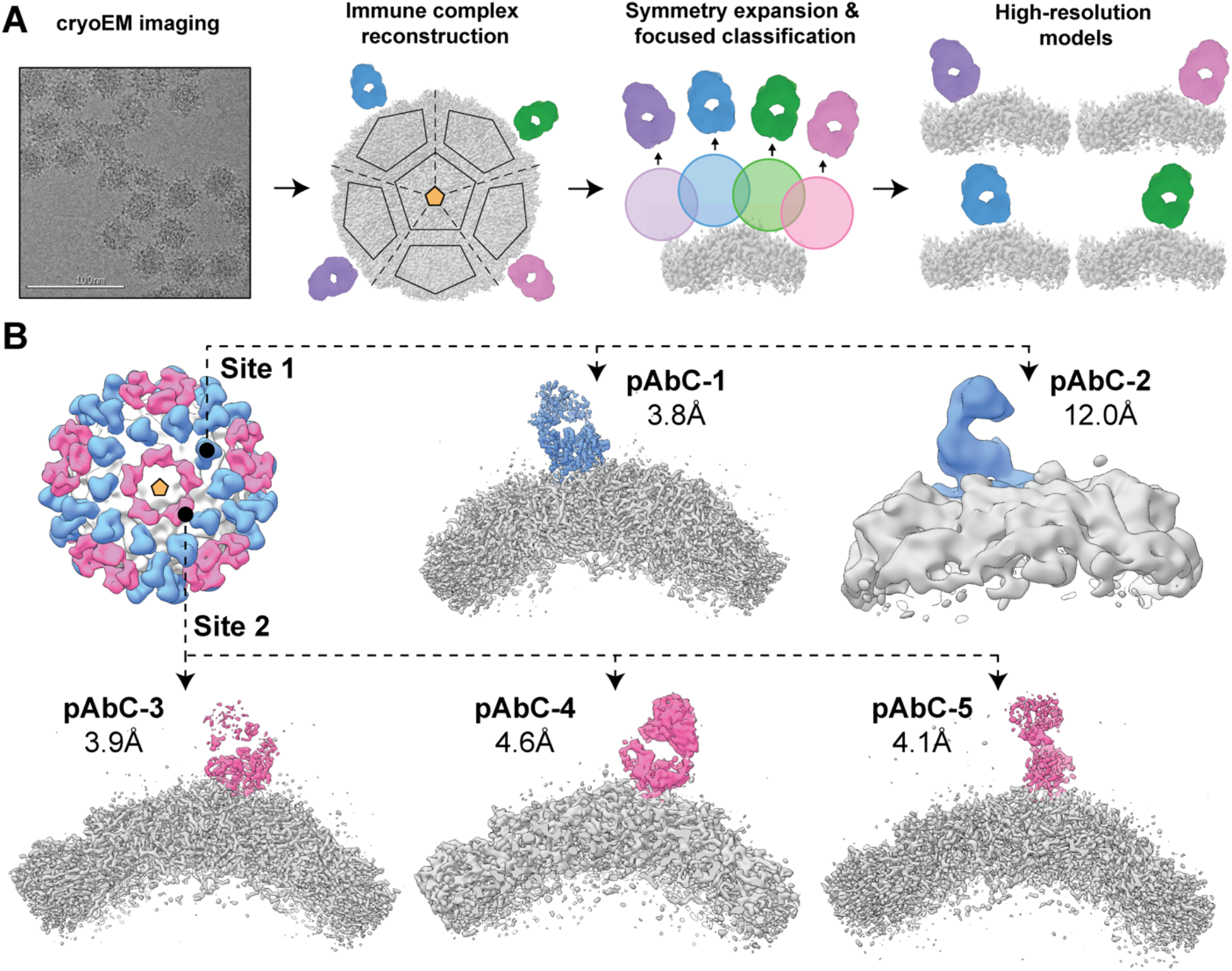
cryoEMPEM analysis of bound polyclonal antibodies. **[A]** Schematic illustration of the cryoEMPEM workflow. **[B]** High resolution antigen-pAb immune complexes featuring structurally unique antibody specificities detected in cryoEMPEM dataset. Antigen is represented in gray and pAb densities are colored according to the epitope (blue and pink colors are used to represent Sites 1 and 2, respectively). The apparent global resolution of each reconstructed cryoEM map is indicated. The most relevant icosahedral symmetry axes are indicated with golden-yellow pentamers (5-fold), triangles (3-fold) and ovals (2-fold).

We positioned spherical solvent masks around 5 epitope clusters (Figure 2 – figure supplement 3A). Antibodies were only detected at Site-1 and Site-2 clusters, consistent with the nsEM data. We resolved two distinct polyclonal antibodies targeting Site-1 and three against Site-2 (Figure 2B, Figure 2 – figure supplement 3B). The highest achieved global map resolution was 3.8Å (pAbC-1, Figure 2 – figure supplement 4A), although the local resolution was worse for the pAb-corresponding part of the map (Figure 2 – figure supplement 4B). Specific epitope contacts and angle of approach were highly similar for Site-1 antibodies while Site-2 pAbs displayed greater diversity (Figure 2 – figure supplement 3B); consistent with nsEM analysis.

We next relaxed atomic models into the pAbC-1, pAbC-3 and pAbC-5 cryoEMPEM maps. Given the polyclonal nature of bound antibodies and the inherent lack of sequence information, the variable Fab domains (Fv) were represented as poly-Alanine models. The models were not built for pAbC-2 and pAbC-4 due to relatively low map resolutions. Instead, we docked mock antibody (PDB ID: 3i9g; (Wojciak et al., 2009)) and previously published CV-A21 structure (PDB ID:1z7s; (Xiao et al., 2005)) into these maps (Figure 3 – figure supplement 1). CryoEMPEM maps and corresponding models were used to assign specific epitope contacts for Site-1 (Figure 3 – figure supplement 2) and Site-2 pAbs (Figure 3 – figure supplement 3).

**Figure 3.**
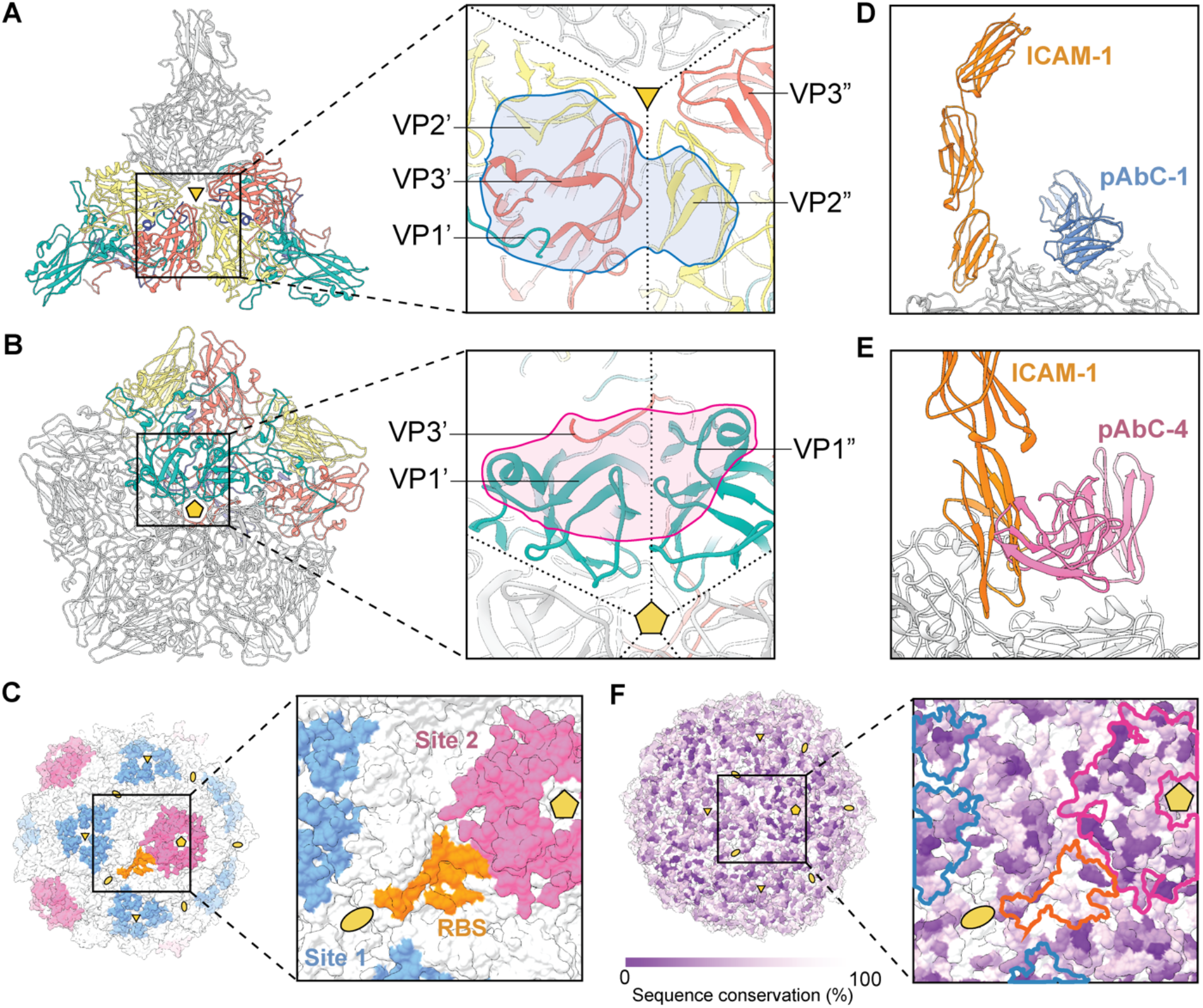
Antibody binding analysis based on cryoEMPEM data. **[A, B]** Locations of the Site-1 (A) and Site-2 (B) epitope clusters. 3 VP protomers are shown for Site-1, and 5 protomers are shown for Site-2. For clarity, epitopes are presented only on a single protomer-protomer interface and the protomers comprising the interface are colored based on the specific VP subunit (VP1 – green; VP2 – yellow; VP3 – salmon; VP4 – dark blue). VP1’ – VP4’ belong to the first protomer in the interface and VP1” – VP4” to the second protomer. Ribbon representation is used in the panel. Polyclonal antibody footprints are indicated in blue (Site-1) and pink (Site-2). **[C]** Surface representation of the CV-A21 viral particle with Site-1 (blue), Site-2 (pink) and the receptor binding site (RBS; orange) indicated. For clarity, only a single RBS is highlighted. **[D,E]** Overlay of pAbC-1 (D) and pAbC-4 (E) antibody structures and ICAM-1 receptor structure (PDB ID: 1z7z; (Xiao et al., 2005)). **[F]** Surface representation of the CV-A21 viral particle colored based on sequence conservation. The location of the Site-1, Site-2 and RBS is outlined in blue, pink and orange, respectively. The most relevant symmetry axes are indicated with golden-yellow pentamers (5-fold), triangles (3-fold) and ovals (2-fold).

Antibodies to Site-1 and Site-2 use different combinations of VP components for binding. Site-1 comprises VP1, VP2 and VP3 elements, while Site-2 is primarily formed by VP1 with small contribution from the C-terminus of VP3 (Figure 3A, Figure 3 – figure supplement 2-4). pAbC-1 and pAbC-5 make contact with residues in both protomers and require a fully assembled viral particle to bind (Figure 3 – figure supplement 2-3). pAbC-2, pAbC-3 and pAbC-4 interact with residues within a single protomer, but from different VP components. Importantly, all pAbs bind in a manner that does not block the accessibility to the symmetry-related neighboring binding site. In theory, all 3 epitopes surrounding the 3-fold symmetry axis and all 5 epitopes at the 5-fold symmetry axis could be simultaneously occupied with antibodies. We speculate that the relatively high immunogenicity of Site-1 ad Site-2 epitope clusters is due to multivalent binding to IgG and IgM forms of antibodies; as previously described for human papillomavirus (Schiller & Lowy, 2018).

Site-2 is particularly interesting because it is in immediate proximity to the receptor binding site (RBS) for CV-A21 (Figure 3C). While antibodies targeting Site-1 are distal from the RBS and it is unlikely that they would interfere with receptor binding as Fabs (Figure 3D), antibodies against Site-2 sterically block ICAM-1 from accessing the RBS (Figure 3E) and may be able to suppress viral entry. We then looked at the sequence conservation across all enterovirus C CV-A strains: A1, A11, A13, A17, A18, A19, A20, A21, A22, A24 (Figure 3F, Figure 3 – figure supplement 5-7). While both Site-1 and Site-2 are highly immunogenic, the sequence conservation is relatively low, and elicited antibody responses are unlikely to cross-react with other strains of CV.

We compared the pAbs identified in this study to the structures of previously isolated mAbs targeting different CV strains (structural data is available for mAb complexes with A6, A10, A16 and B1 viruses). Overall, we observed a high degree of overall similarity in epitopes for several of the antibodies. For example, Site-1 overlaps with the binding sites of 14B10 (He et al., 2020), 5F5 (Xu et al., 2021) and 2G8 (Zhu et al., 2018) mAbs; the overlay of pAbC-1 and 14B10 mAb structures is shown in Figure 3 – figure supplement 8A. Site-2 pAbs display high similarity in epitope and angle of approach to 1D5 mAb (Xu et al., 2017), and a partial overlap with 18A7 mAb (He et al., 2020) that binds directly on top of the 5-fold symmetry axis (Figure 3 – figure supplement 8B). These findings suggest that there may be a level of plasticity in antibody responses to CV with respect to the locations of immunodominant epitopes.

## Discussion

Here we introduce a method for analysis of polyclonal antibody responses elicited by non-enveloped viruses with an icosahedral capsid. We validated the approach on the coxsackievirus A21 using polyclonal antibodies from mice. In non-enveloped icosahedral virions antibody epitopes are likely to be comprised of amino acids from multiple protomers and/or subunits. Thus, intact viral particles with native antigen presentation, instead of minimal subunit antigens, are necessary for comprehensive evaluation of elicited antibody responses. Our EMPEM methods can now be readily implemented to other non-enveloped icosahedral viruses (e.g., papillomaviruses, parvoviruses, adenoviruses), as well as enveloped viruses with icosahedral lattice on their surface (e.g., flaviviruses, togaviruses).

From a practical standpoint, the application of high-symmetry viral particles with relatively small diameter (~30 nm) allows one to generate relatively large particle subsets after symmetry expansion (by a factor of 60). For comparison, while the diameter of homotrimeric glycoprotein spikes from flu (HA), HIV (Env) and SARS-CoV-2 (S) is ~10-15 nm, the increase in particle subsets after symmetry expansion is only 3-fold. Larger starting datasets increase the number of potential pAb-epitope combinations that can be analyzed in parallel, improve the chances of finding low abundance pAbs, and increase the resolution that can be achieved.

Using a combination of nsEM- and cryoEM-based polyclonal epitope mapping we identified two primary immunogenic sites on the surface of CV-A21 located near the 3-fold and 5-fold axes. All reconstructed antibodies bind in a manner that allows equivalent neighboring sites on the icosahedral lattice to be simultaneously occupied by the same type of antibody and achieve maximum stoichiometry of 60:1 (Fab to virus). Average distance between symmetry-related epitopes is ~30-60 Å which is comparable to HPV where epitopes are spaced by ~50-100 Å (Schiller & Lowy, 2018). Such ordered array of epitopes allows for multivalent binding by antigen-specific B-cell receptors resulting in strong activation of the downstream signaling cascade in these B cells. This is the likely explanation for robust antibody response against epitopes proximal to the symmetry axis but the lack of antibodies directly on the symmetry axis. Consistently, the majority of isolated monoclonal antibodies against different coxsackievirus strains (for which structural data exist) target similar epitopes. The exceptions are 18A7 mAb with an epitope directly on the 5-fold symmetry axis and NA9D7 mAb interacting with the receptor binding site (He et al., 2020). Using EMPEM we can study polyclonal antibody responses in a longitudinal manner enabling us to understand the hierarchy of immunodominant epitopes in different CV strains and to identify optimal vaccine candidates for eliciting the most desirable antibody responses.

Antibodies targeting epitopes near the 5-fold axis can sterically block access to the receptor binding site and thereby prevent the virus from infecting cells. This feature is highly desirable, but the relatively low level of sequence conservation in this area is problematic as elicited antibodies are unlikely to cross-react with other coxsackievirus strains and may sterically compete with more desirable antibodies. Traditionally, this is achieved by immunogen engineering with many examples from the RSV, flu, HIV and SARS-CoV-2 fields, or through viral cocktail immunizations as done with HPV and Dengue vaccines (Bangaru et al., 2020; Cheng, Wang, & Du, 2020; Ellis et al., 2021; McLellan et al., 2013; Medina-Ramirez et al., 2017; Ou et al., 2020; Steichen et al., 2019; Wei et al., 2020; Wilder-Smith, 2020). Similar vaccine design strategies can be employed to coxsackieviruses and EM methods presented in this study open the door for rapid assessment of new vaccine candidates.

## Materials and Methods

### Preparation of Coxsackievirus A21 viral particles

Coxsackievirus A21 particles were propagated in H1-HeLa cells. The virions were harvested from lysed cells 2 days after infection, clarified by centrifugation, 0.2 μm filtered, and inactivated by incubation with 100 μg/mL formaldehyde at 37 °C for 3 days, followed by low-speed centrifugation to remove cell debris. The supernatant was pelleted through a 30% sucrose cushion via ultracentrifugation at 175,000 g for 14 hours at 4 °C. The resuspended pellet was sedimented through a 15-45% (w/v) sucrose density gradient and centrifuged at 120,000 g for 14 hours at 4 °C. Fractions containing virus particles were collected and dialyzed against PBS buffer. Inactivated particles were characterized and used for EMPEM and high-resolution structural analysis using cryo-EM. The efficacy of formaldehyde treatment on virus infection was evaluated by the infectivity assay visualizing cytopathic effects in comparison to untreated controls. Total protein in virus fractions was determined and the presence of viral proteins visualized by Western blot. For structural studies, the gradient purified viral particles were polished by size-exclusion chromatography using the HiPrep 16/60 Sephacryl S-500 HR column (Sigma Aldrich) running in TBS buffer (Alfa Aesar). Fractions corresponding to Coxsackievirus A21 particles were pooled and concentrated to 0.5mg/ml using Amicon Ultra centrifugal filter units with 100 kDa cutoff (Millipore Sigma).

### Mouse immunization experiments

Animal experiments were performed at Mayo Clinic with permissions from the Mayo Clinic Institutional Animal Care and Use Committee with the registration number A00005024. C57Bl/6 mice were injected intravenously with PBS (n=6) or 1×10^6^ TCID_50_ of purified CV-A21 virus on days 0 and 14 (n=20). A serum sample was collected to verify the presence of neutralizing antibodies prior to euthanizing all mice. At the time of euthanasia (week 4), serum was collected from all immunized mice, pooled, heat-inactivated at 56 °C for 30 minutes and stored at −80 °C. Immunologically naïve mouse serum samples were obtained and processed similarly.

### Preparation of polyclonal antibody samples

~3 mL of serum from CV-A21-immunized or naïve mice was diluted in 97 mL of PBS and ran over a column pre-packed with CaptureSelect IgG-Fc (Multispecies) Affinity Matrix (ThermoFisher Scientific). IgGs were eluted with 0.1 M glycine buffer (pH 3.0) and immediately neutralized with 1 M Tris-HCl pH 8. IgG samples were concentrated and buffer-exchanged to digestion buffer (PBS + 10 mM EDTA + 10 mM cysteine, pH 7.4) using Amicon Ultra centrifugal filter units with 10 kDa cutoff (Millipore Sigma). The digestion into Fab/Fc fragments was performed for 5 hours at 37 °C using 40 μg of activated papain per 1 mg of IgG. Non-digested IgG was removed by size-exclusion chromatography, using the HiLoad 16/600 Superdex 200 pg column (GE Healthcare) running in TBS buffer. Fractions corresponding to the Fab/Fc fragments were pooled and concentrated using Amicon Ultra centrifugal filter units with 10 kDa cutoff. Final Fab/Fc yields were ~10 mg from the starting ~3 mL of serum.

### Preparation of immune complexes

For negative stain EM experiments, we prepared the immune complexes by combining 50 μg of purified, inactivated CV-A21 viral particles with (1) TBS, (2) 500 μg of Fab/Fc from the immunologically naïve mice or (3) 500 μg of Fab/Fc from the mice immunized with CV-A21. The assembly reactions were incubated at room temperature for ~18 hours, and then the samples were subjected to size-exclusion chromatography using Superose 6 Increase 10/300 GL column running in TBS buffer. Fractions corresponding to the CV-A21 virions (or immune complexes with pAbs) were pooled and concentrated using Amicon Ultra centrifugal filter units with 100 kDa cutoff.

For cryoEM the immune complexes were assembled by combining 300 μg of the purified, inactivated CV-A21 viral particles with 8 mg of Fab/Fc from the mice that received CV-A21 virus as immunogen. Further processing was done as described in the previous paragraph.

### Negative stain EM – Grid preparation, Imaging and Data Processing

Samples featuring CV-A21 particles (free or as part of an immune complex) were diluted in TBS to 50 μg/ml and loaded onto in-house made carbon-coated 400-mesh copper grids (glow-discharged at 15 mA for 25 s). The sample was blotted off after 10 s and the grids were stained with 2% (w/v) uranyl-formate for 60 s. Grids were imaged on a Tecnai F20 electron microscope (FEI) equipped with a TemCam F416 CMOS camera (TVIPS). The microscope operates at 200 kV. The imaging defocus was set to 1.5 μm and the total electron dose adjusted to 25 e^−^/Å^2^. The magnification was set to 62,000 X, with the resulting pixel size of 1.77 Å. Leginon software (Suloway et al., 2005) was used for image acquisition and all the early processing steps were performed in Appion (Lander et al., 2009). 2D/3D classification and 3D refinement steps were done in Relion/3.0 (Zivanov et al., 2018). Icosahedral symmetry was imposed for the 3D classification and refinement steps. Negative stain EM maps have been deposited to the EM Data Bank under accession ID: EMD-26065.

### Cryo-EM – Grid preparation

Immune complexes comprising inactivated CV-A21 viral particles and CV-A21-elicited polyclonal antibodies were concentrated to 2.5 mg/ml using Amicon Ultra centrifugal filter units with 100 kDa cutoff (Millipore Sigma). Cryo-grids were prepared on a Vitrobot mark IV (Thermo Fisher Scientific). Vitrobot settings were: Temperature = 10 °C; Humidity = 100%; Blotting force = 0; Wait time = 10 s; Blotting time varied in the 3-6 s range. Quantifoil R 2/1 holey carbon copper grid (EMS) were used for sample vitrification. Prior to sample application the grids were subjected to plasma cleaning (Ar/O2 gas mixture; Solarus 950 plasma cleaner, Gatan). 3 μL of the sample was loaded onto plasma-cleaned grid on Vitrobot. Following the blotting step, the grids were plunged into liquid-nitrogen-cooled liquid ethane.

### Cryo-EM – Imaging and Data Processing

Samples were imaged on FEI Titan Krios electron microscope (ThermoFisher) operating at 300 kV. The microscope was equipped with the K2 summit detector (Gatan) running in counting mode. Nominal magnification was set to 130,000 X with the pixel size of 1.045 Å (at the specimen plane). Automated data collection was performed using Leginon (Suloway et al., 2005). Data collection information is provided in Figure 2 – figure supplement 1. Raw micrograph frames were aligned and dose-weighted using MotionCor2 (Zheng et al., 2017). Micrographs were then imported to Relion/3.0 for further processing (Zivanov et al., 2018). CTF estimation was performed using GCTF (Zhang, 2016). Particles were picked manually and subjected to 1 round of 2D classification to remove bad picks. Selected particles were subjected to a round of 3D refinement with icosahedral symmetry imposed. Low resolution map of the CV capsid was used as an initial model for the initial 3D refinement/classification steps. A tight solvent mask around the CV capsid was applied to remove the signal contributions from pAbs and internal viral components during refinement. Pre-refined particles were symmetry-expanded (icosahedral symmetry). This increased the size of the dataset by 60-fold and collapsed all bound pAbs onto every symmetry-related protomer, thus facilitating the classification of different pAbs. To avoid the possibility of symmetry-related copies from aligning to themselves we restricted the movement of particles during the subsequent 3D refinement and classification steps. Particle alignment was suspended for 3D classification (--skip_align, T 16), and only local angular searches were allowed for 3D refinement (1.8° angular step per iteration, --healpix_order 4 -- auto_local_healpix_order 4). For classification, spherical masks (80 Å diameter) were positioned around different potential pAb epitopes on a single protomer (2 spots) as well as directly on the 2-fold, 3-fold and 5-fold symmetry axes. In the 1^st^ round, separate 3D classifications were performed for each of the 6 masks used (sorted into 40 3D classes). pAb-containing classes with unique structural features (i.e., epitope and orientation) were selected and treated independently after this step. For each of the selected particle subsets, we extracted subparticle regions centering on the pAb. This was done to reduce the alignment bias from the CV particle during 3D refinement; the molecular weight of CV capsid is ~6 MDa while Fabs are only 50 kDa. pAb-centered subparticles were subjected to a round of 3D refinement with solvent mask around the immune complex and restricted angular search space (local angular searches only). Following refinement, each subset of particles was then subjected to 1 round of 3D classification with a spherical mask (80 Å diameter) centered on each respective pAb (--skip_align, T 16, K 5). The highest quality classes based on structural features and estimated resolution were selected for the next round of classification. Here, 3D classification was performed with a solvent mask surrounding the entire subparticle comprising pAb and part of CV capsid (--skip_align, T 16, K 3). The highest quality classes based on structural features and estimated resolution were selected and subjected to 3D refinement (same settings as in the previous refinement step). The maps from 3D refinement were Postprocessed in Relion. Postprocessing consisted of B-factor sharpening, MTF correction and solvent masking. The resulting maps were used for model building and submission to EM Data Bank (accession IDs: EMD-26072, EMD-26069, EMD-26068, EMD-26070, EMD-26071). Data processing workflows and relevant information are presented in Figure 2 – figure supplement 2. Data processing information are presented in Figure 3 – figure supplement 1.

### Model building and refinement

Atomic models were built into postprocessed maps from Relion. CV-A21 structure from PDB entry 1z7s (Xiao et al., 2005) was used as an initial model for the antigen. Only the protomers proximal to the pAb epitope were built into the reconstructed maps. Initial pAb model (Fv-domain only) were generated from PDB entry 3i9g (Wojciak et al., 2009) by mutating all of the amino acid residues to alanine. UCSF Chimera (Pettersen et al., 2004) was used to dock the antigens/pAbs into the reconstructed maps and create the initial models. The models were then relaxed by iterating manual refinement steps in Coot (Emsley & Crispin, 2018) and automated refinement steps in Rosetta (Wang et al., 2016). The number of residues in each pAb model was adjusted to best recapitulate the structural features within each cryoEMPEM map. Structural homology to published mouse antibody structures (primarily 3i9g (Wojciak et al., 2009)) was used to assign antibody heavy (H) and light (L) chains and individual CDR loops. The principal factors for discrimination are described previously (Antanasijevic et al., 2021). MolProbity (Williams et al., 2018) and EMRinger (Barad et al., 2015) metrics, were used to assess model quality. The final statistics are shown in Figure 3 – figure supplement 1. All models were submitted to the Protein Data Bank (accession IDs: 7TQU, 7TQS, 7TQT).

## Acknowledgements

The authors acknowledge Bill Anderson, Hannah L. Turner, Charles A. Bowman and Jean-Christophe Ducom (The Scripps Research Institute) for their help with different aspects of EM experiments. The authors thank Lauren Holden for her help on the preparation of this manuscript. We also sincerely and respectfully acknowledge the revolutionary contribution to medical research made possible by Henrietta Lacks and HeLa cells.

## Author contributions

A.B.W., V.S.R., A.J.S., and A.A. conceived and supervised the study. A.J.S. and V.S.R. prepared the CV-A21 antigen, designed and executed the mouse immunization experiments. A.A. purified polyclonal antibodies and produced the immune complexes. A.A. performed all EM experiments and processed the data. A.A., V.S.R., and A.B.W. wrote the initial draft of the manuscript. All authors contributed to the final manuscript text by assisting in writing and providing critical feedback.

## Data and materials availability

All data needed to evaluate the conclusions in the paper are present in the manuscript and/or the Supplementary Materials. 3D maps have been deposited to the Electron Microscopy Data Bank (http://www.emdatabank.org/) with accession numbers: EMD-26065, EMD-26072, EMD-26069, EMD-26068, EMD-26070, EMD-26071. Atomic models have been deposited to the Protein Data Bank (http://www.rcsb.org/) with accession numbers: 7TQU, 7TQS, 7TQT.

## References

Antanasijevic, A., Bowman, C. A., Kirchdoerfer, R. N., Cottrell, C. A., Ozorowski, G., Upadhyay, A. A., … Ward, A. B. (2022). From structure to sequence: Antibody discovery using cryoEM. Sci Adv, 8(3), eabk2039. doi:10.1126/sciadv.abk2039

Antanasijevic, A., Sewall, L. M., Cottrell, C. A., Carnathan, D. G., Jimenez, L. E., Ngo, J. T., … Ward, A. B. (2021). Polyclonal antibody responses to HIV Env immunogens resolved using cryoEM. Nat Commun, 12(4817). doi:https://doi.org/10.1038/s41467-021-25087-4

Bangaru, S., Antanasijevic, A., Kose, N., Sewall, L. M., Jackson, A. M., Suryadevara, N., … Ward, A. B. (2021). Structural mapping of antibody landscapes to human betacoronavirus spike proteins. bioRxiv. doi:https://doi.org/10.1101/2021.09.30.462459

Bangaru, S., Ozorowski, G., Turner, H. L., Antanasijevic, A., Huang, D., Wang, X., … Ward, A. B. (2020). Structural analysis of full-length SARS-CoV-2 spike protein from an advanced vaccine candidate. Science. doi:10.1126/science.abe1502

Barad, B. A., Echols, N., Wang, R. Y., Cheng, Y., DiMaio, F., Adams, P. D., & Fraser, J. S. (2015). EMRinger: side chain-directed model and map validation for 3D cryo-EM. Nat Methods, 12(10), 943-946. doi:10.1038/nmeth.3541

Basavappa, R., Syed, R., Flore, O., Icenogle, J. P., Filman, D. J., & Hogle, J. M. (1994). Role and mechanism of the maturation cleavage of VP0 in poliovirus assembly: structure of the empty capsid assembly intermediate at 2.9 A resolution. Protein Sci, 3(10), 1651-1669. doi:10.1002/pro.5560031005

Bianchi, M., Turner, H. L., Nogal, B., Cottrell, C. A., Oyen, D., Pauthner, M., … Hangartner, L. (2018). Electron-Microscopy-Based Epitope Mapping Defines Specificities of Polyclonal Antibodies Elicited during HIV-1 BG505 Envelope Trimer Immunization. Immunity, 49(2), 288-300 e288. doi:10.1016/j.immuni.2018.07.009

Bradley, S., Jakes, A. D., Harrington, K., Pandha, H., Melcher, A., & Errington-Mais, F. (2014). Applications of coxsackievirus A21 in oncology. Oncolytic Virother, 3, 47-55. doi:10.2147/OV.S56322

Cheng, L., Wang, Y., & Du, J. (2020). Human Papillomavirus Vaccines: An Updated Review. Vaccines (Basel), 8(3). doi:10.3390/vaccines8030391

Couch, R. B., Douglas, R. G., Jr., Lindgren, K. M., Gerone, P. J., & Knight, V. (1970). Airborne transmission of respiratory infection with coxsackievirus A type 21. Am J Epidemiol, 91(1), 78-86. doi:10.1093/oxfordjournals.aje.a121115

Curry, S., Fry, E., Blakemore, W., Abu-Ghazaleh, R., Jackson, T., King, A., … Stuart, D. (1997). Dissecting the roles of VP0 cleavage and RNA packaging in picornavirus capsid stabilization: the structure of empty capsids of foot-and-mouth disease virus. J Virol, 71(12), 9743-9752. doi:10.1128/JVI.71.12.9743-9752.1997

Ellis, D., Brunette, N., Crawford, K. H. D., Walls, A. C., Pham, M. N., Chen, C., … King, N. P. (2021). Stabilization of the SARS-CoV-2 Spike Receptor-Binding Domain Using Deep Mutational Scanning and Structure-Based Design. Front Immunol, 12, 710263. doi:10.3389/fimmu.2021.710263

Emsley, P., & Crispin, M. (2018). Structural analysis of glycoproteins: building N-linked glycans with Coot. Acta Crystallogr D Struct Biol, 74(Pt 4), 256-263. doi:10.1107/S2059798318005119

Granoff, A., & Webster, R. G. (1999). Encyclopedia of virology (2nd ed.). San Diego: Academic Press.

Guthmiller, J. J., Han, J., Utset, H. A., Li, L., Lan, L. Y., Henry, C., … Wilson, P. C. (2021). Broadly neutralizing antibodies target a hemagglutinin anchor epitope. Nature. doi:10.1038/s41586-021-04356-8

He, M., Xu, L., Zheng, Q., Zhu, R., Yin, Z., Zha, Z., … Xia, N. (2020). Identification of Antibodies with Non-overlapping Neutralization Sites that Target Coxsackievirus A16. Cell Host Microbe, 27(2), 249-261 e245. doi:10.1016/j.chom.2020.01.003

Lander, G. C., Stagg, S. M., Voss, N. R., Cheng, A., Fellmann, D., Pulokas, J., … Carragher, B. (2009). Appion: an integrated, database-driven pipeline to facilitate EM image processing. J Struct Biol, 166(1), 95-102.

Li, N., Li, Z., Fu, Y., & Cao, S. (2020). Cryo-EM Studies of Virus-Antibody Immune Complexes. Virol Sin, 35(1), 1-13. doi:10.1007/s12250-019-00190-5

McCarthy, C., Jayawardena, N., Burga, L. N., & Bostina, M. (2019). Developing Picornaviruses for Cancer Therapy. Cancers (Basel), 11(5). doi:10.3390/cancers11050685

McLellan, J. S., Chen, M., Joyce, M. G., Sastry, M., Stewart-Jones, G. B., Yang, Y., … Kwong, P. D. (2013). Structure-based design of a fusion glycoprotein vaccine for respiratory syncytial virus. Science, 342(6158), 592-598. doi:10.1126/science.1243283

Medina-Ramirez, M., Garces, F., Escolano, A., Skog, P., de Taeye, S. W., Del Moral-Sanchez, I., … Sanders, R. W. (2017). Design and crystal structure of a native-like HIV-1 envelope trimer that engages multiple broadly neutralizing antibody precursors in vivo. J Exp Med, 214(9), 2573-2590. doi:10.1084/jem.20161160

Murphy, O. C., Messacar, K., Benson, L., Bove, R., Carpenter, J. L., Crawford, T., … group, A. F. M. w. (2021). Acute flaccid myelitis: cause, diagnosis, and management. Lancet, 397(10271), 334-346. doi:10.1016/S0140-6736(20)32723-9

Nikonov, O. S., Chernykh, E. S., Garber, M. B., & Nikonova, E. Y. (2017). Enteroviruses: Classification, Diseases They Cause, and Approaches to Development of Antiviral Drugs. Biochemistry (Mosc), 82(13), 1615-1631. doi:10.1134/S0006297917130041

Nogal, B., Bianchi, M., Cottrell, C. A., Kirchdoerfer, R. N., Sewall, L. M., Turner, H. L., … Ward, A. B. (2020). Mapping Polyclonal Antibody Responses in Non-human Primates Vaccinated with HIV Env Trimer Subunit Vaccines. Cell Rep, 30(11), 3755-3765 e3757. doi:10.1016/j.celrep.2020.02.061

Ou, L., Kong, W. P., Chuang, G. Y., Ghosh, M., Gulla, K., O’Dell, S., … Kwong, P. D. (2020). Preclinical Development of a Fusion Peptide Conjugate as an HIV Vaccine Immunogen. Sci Rep, 10(1), 3032. doi:10.1038/s41598-020-59711-y

Oyero, O. G., Adu, F. D., & Ayukekbong, J. A. (2014). Molecular characterization of diverse species enterovirus-B types from children with acute flaccid paralysis and asymptomatic children in Nigeria. Virus Res, 189, 189-193. doi:10.1016/j.virusres.2014.05.029

Pettersen, E. F., Goddard, T. D., Huang, C. C., Couch, G. S., Greenblatt, D. M., Meng, E. C., & Ferrin, T. E. (2004). UCSF Chimera--a visualization system for exploratory research and analysis. J Comput Chem, 25(13), 1605-1612. doi:10.1002/jcc.20084

Puenpa, J., Wanlapakorn, N., Vongpunsawad, S., & Poovorawan, Y. (2019). The History of Enterovirus A71 Outbreaks and Molecular Epidemiology in the Asia-Pacific Region. J Biomed Sci, 26(1), 75. doi:10.1186/s12929-019-0573-2

Schiller, J., & Lowy, D. (2018). Explanations for the high potency of HPV prophylactic vaccines. Vaccine, 36(32 Pt A), 4768-4773. doi:10.1016/j.vaccine.2017.12.079

Stass, R., Ilca, S. L., & Huiskonen, J. T. (2018). Beyond structures of highly symmetric purified viral capsids by cryo-EM. Curr Opin Struct Biol, 52, 25-31. doi:10.1016/j.sbi.2018.07.011

Steichen, J. M., Lin, Y. C., Havenar-Daughton, C., Pecetta, S., Ozorowski, G., Willis, J. R., … Schief, W. R. (2019). A generalized HIV vaccine design strategy for priming of broadly neutralizing antibody responses. Science, 366(6470). doi:10.1126/science.aax4380

Suloway, C., Pulokas, J., Fellmann, D., Cheng, A., Guerra, F., Quispe, J., … Carragher, B. (2005). Automated molecular microscopy: the new Leginon system. J Struct Biol, 151(1), 41-60. doi:10.1016/j.jsb.2005.03.010

Wang, R. Y., Song, Y., Barad, B. A., Cheng, Y., Fraser, J. S., & DiMaio, F. (2016). Automated structure refinement of macromolecular assemblies from cryo-EM maps using Rosetta. Elife, 5. doi:10.7554/eLife.17219

Ward, A. B., & Wilson, I. A. (2017). The HIV-1 envelope glycoprotein structure: nailing down a moving target. Immunol Rev, 275(1), 21-32. doi:10.1111/imr.12507

Wei, C. J., Crank, M. C., Shiver, J., Graham, B. S., Mascola, J. R., & Nabel, G. J. (2020). Next-generation influenza vaccines: opportunities and challenges. Nat Rev Drug Discov, 19(4), 239-252. doi:10.1038/s41573-019-0056-x

Wilder-Smith, A. (2020). Dengue vaccine development: status and future. Bundesgesundheitsblatt Gesundheitsforschung Gesundheitsschutz, 63(1), 40-44. doi:10.1007/s00103-019-03060-3

Williams, C. J., Headd, J. J., Moriarty, N. W., Prisant, M. G., Videau, L. L., Deis, L. N., … Richardson, D. C. (2018). MolProbity: More and better reference data for improved all-atom structure validation. Protein Sci, 27(1), 293-315. doi:10.1002/pro.3330

Wojciak, J. M., Zhu, N., Schuerenberg, K. T., Moreno, K., Shestowsky, W. S., Hiraiwa, M., … Huxford, T. (2009). The crystal structure of sphingosine-1-phosphate in complex with a Fab fragment reveals metal bridging of an antibody and its antigen. Proc Natl Acad Sci U S A, 106(42), 17717-17722. doi:10.1073/pnas.0906153106

Xiao, C., Bator-Kelly, C. M., Rieder, E., Chipman, P. R., Craig, A., Kuhn, R. J., … Rossmann, M. G. (2005). The crystal structure of coxsackievirus A21 and its interaction with ICAM-1. Structure, 13(7), 1019-1033. doi:10.1016/j.str.2005.04.011

Xu, L., Zheng, Q., Li, S., He, M., Wu, Y., Li, Y., … Xia, N. (2017). Atomic structures of Coxsackievirus A6 and its complex with a neutralizing antibody. Nat Commun, 8(1), 505. doi:10.1038/s41467-017-00477-9

Xu, L., Zheng, Q., Zhu, R., Yin, Z., Yu, H., Lin, Y., … Xia, N. (2021). Cryo-EM structures reveal the molecular basis of receptor-initiated coxsackievirus uncoating. Cell Host Microbe, 29(3), 448-462 e445. doi:10.1016/j.chom.2021.01.001

Zhang, K. (2016). Gctf: Real-time CTF determination and correction. J Struct Biol, 193(1), 1-12. doi:10.1016/j.jsb.2015.11.003

Zheng, S. Q., Palovcak, E., Armache, J. P., Verba, K. A., Cheng, Y., & Agard, D. A. (2017). MotionCor2: anisotropic correction of beam-induced motion for improved cryo-EM. Nat Methods, 14(4), 331-332. doi:10.1038/nmeth.4193

Zhu, R., Xu, L., Zheng, Q., Cui, Y., Li, S., He, M., … Xia, N. (2018). Discovery and structural characterization of a therapeutic antibody against coxsackievirus A10. Sci Adv, 4(9), eaat7459. doi:10.1126/sciadv.aat7459

Zivanov, J., Nakane, T., Forsberg, B. O., Kimanius, D., Hagen, W. J., Lindahl, E., & Scheres, S. H. (2018). New tools for automated high-resolution cryo-EM structure determination in RELION-3. Elife, 7. doi:10.7554/eLife.42166

Zou, L., Yi, L., Song, Y., Zhang, X., Liang, L., Ni, H., … Lu, J. (2017). A cluster of coxsackievirus A21 associated acute respiratory illness: the evidence of efficient transmission of CVA21. Arch Virol, 162(4), 1057-1059. doi:10.1007/s00705-016-3201-4

