## Supplementary Information for "High-resolution structural analysis of enterovirus-reactive polyclonal antibodies in complex with whole virions"

**Title:**

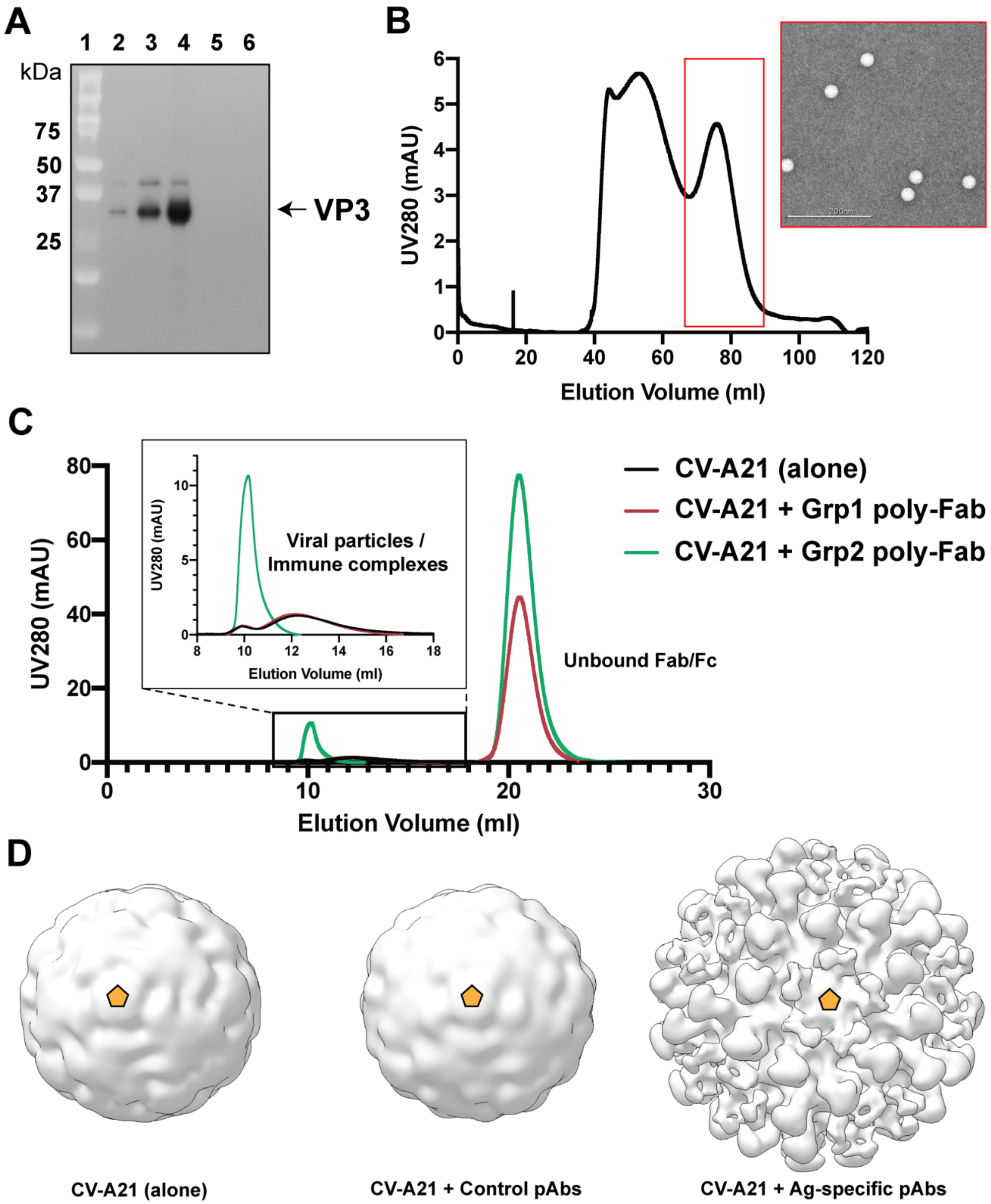


**Figure 1 - figure supplement 1. Purification and nsEM characterization of CV-A21 viruses and immune complexes.** **[A]** Western blot obtained using an Enterovirus pan monoclonal antibody L66J (ThermoFisher; MA5-18206); Lane 1: protein ladder; Lane 2: unconcentrated virus; Lane 3: 30% sucrose purified virus; Lane 4: gradient purified virus; Lane 5: PBS gradient purification control; Lane 6: H1-HeLa cell lysate control. **[B]** SEC purification of formaldehyde-treated CV-A21 particles. Sephacryl S-500 HR column was used for the purification step. Elution peak and nsEM micrographs corresponding to CV-A21 are shown in red. **[C]** SEC purification of CV-A21 viral particles complexed with pAb samples from immunized mice (green and red lines). Noncomplexed CV-A21was also ran as a control (black line). Superose 6 increase 10/300 gl column was used for the SEC step. [D] 3D maps of CV-A21-containing immune complexes reconstructed with icosahedral symmetry imposed. Forward-most facing 5-fold symmetry axis is indicated using golden-yellow pentamer.

**Figure 2 – figure supplement 1. Cryo-EM data collection information**

| | **CV-A21 + Ag-specific mouse pAb** |
|---|---|
| **Microscope** | Titan Krios |
| **Voltage (kV)** | 300 |
| **Detector** | Gatan K2 Summit |
| **Recording mode** | Counting |
| **Magnification** | 130,000 |
| **Movie micrograph pixel size** | 1.045 |
| **Dose rate ($e^{-}/Å^{2}/s$)** | 5.55 |
| **No. of frames per movie micrograph** | 45 |
| **Frame exposure time (ms)** | 200 |
| **Movie micrograph exposure time (s)** | 9.0 |
| **Total dose ($e^{-}/Å^{2}$)** | 49.95 |
| **Grid Type** | Quantifoil R 2/1 |
| **Under focus range (μm)** | 0.8 – 1.8 |
| **Number of movie micrographs** | 3,862 |

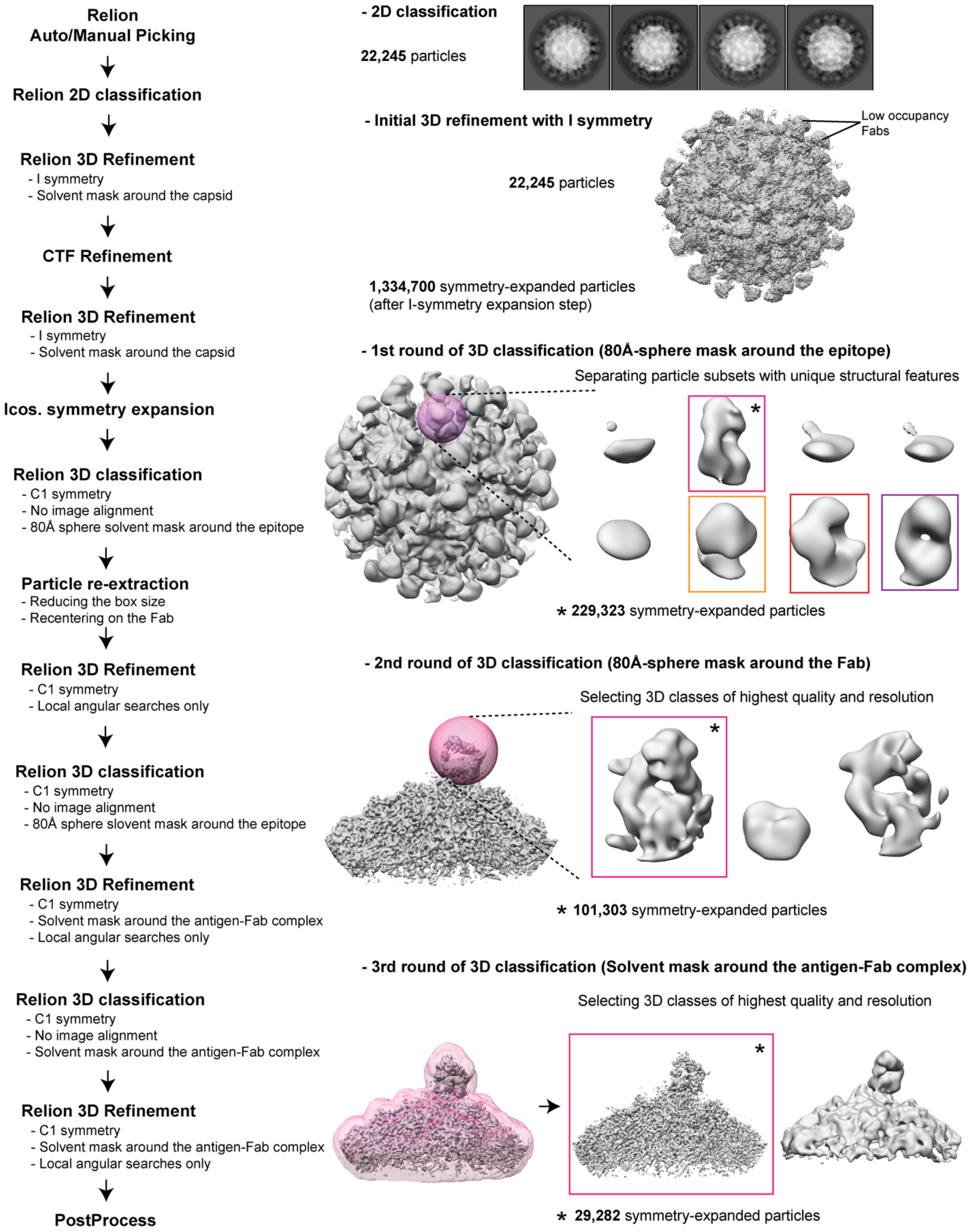


**Figure 2 – figure supplement 2. Schematic representation of the focused classification approach used for processing of cryoEMPEM data.** Full data processing workflow is shown on the left and the examples of intermediate results are shown on the right.

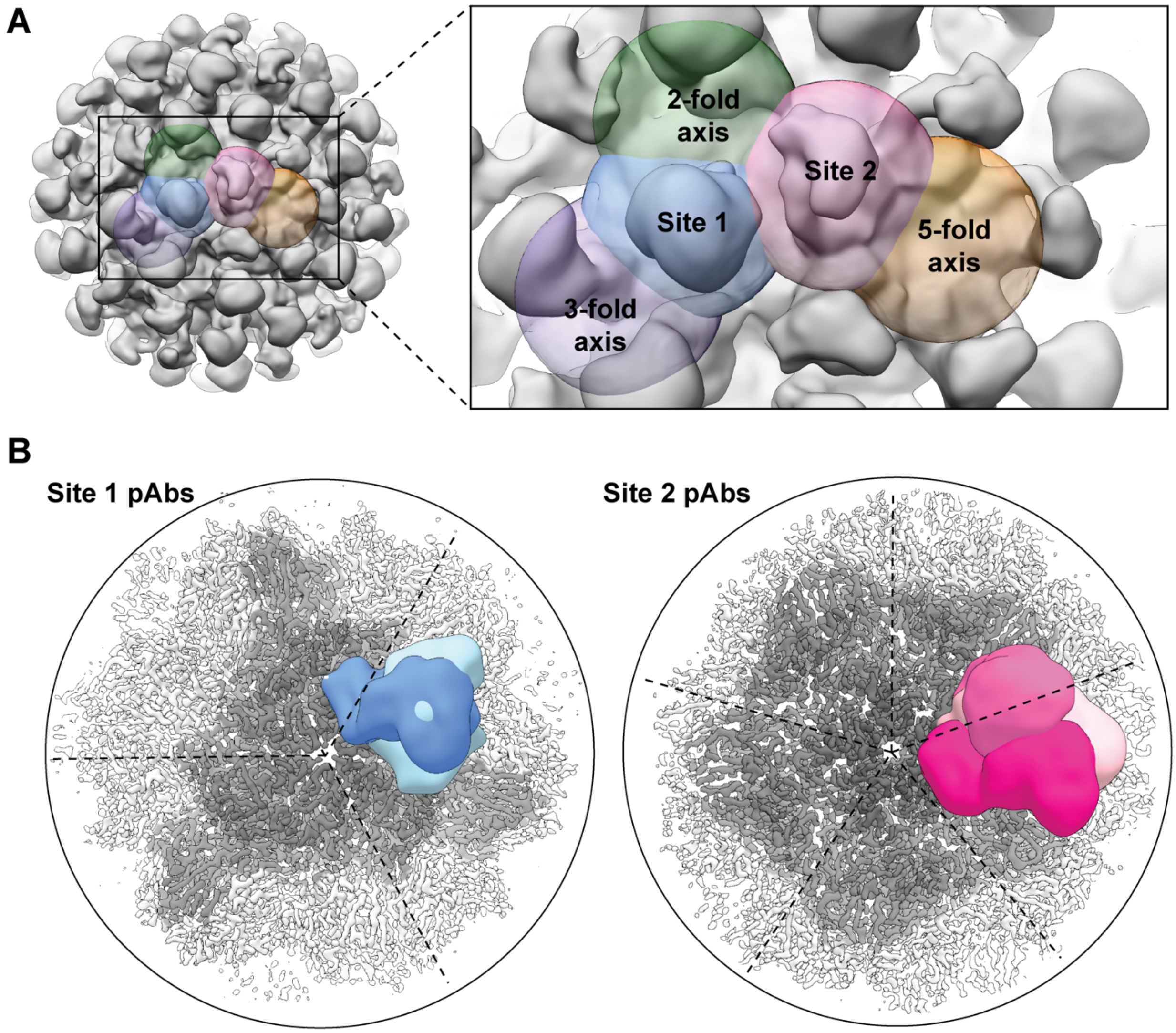


**Figure 2 – figure supplement 3. Solvent masks used for data processing and overlay of reconstructed maps. [A]** The positions of spherical solvent masks used for focused classification of antigen-bound pAbs. **[B]** Overlay of the reconstructed pAbs at Site 1 (left) and Site 2 (right). Different pAbs are shown in different shades of blue (Site 1) and pink (Site 2). pAb-corresponding density in each overlayed map was low-pass filtered to 15Å resolution for clarity. The location of the 3-fold and the 5-fold symmetry axes are indicated in the corresponding panel.

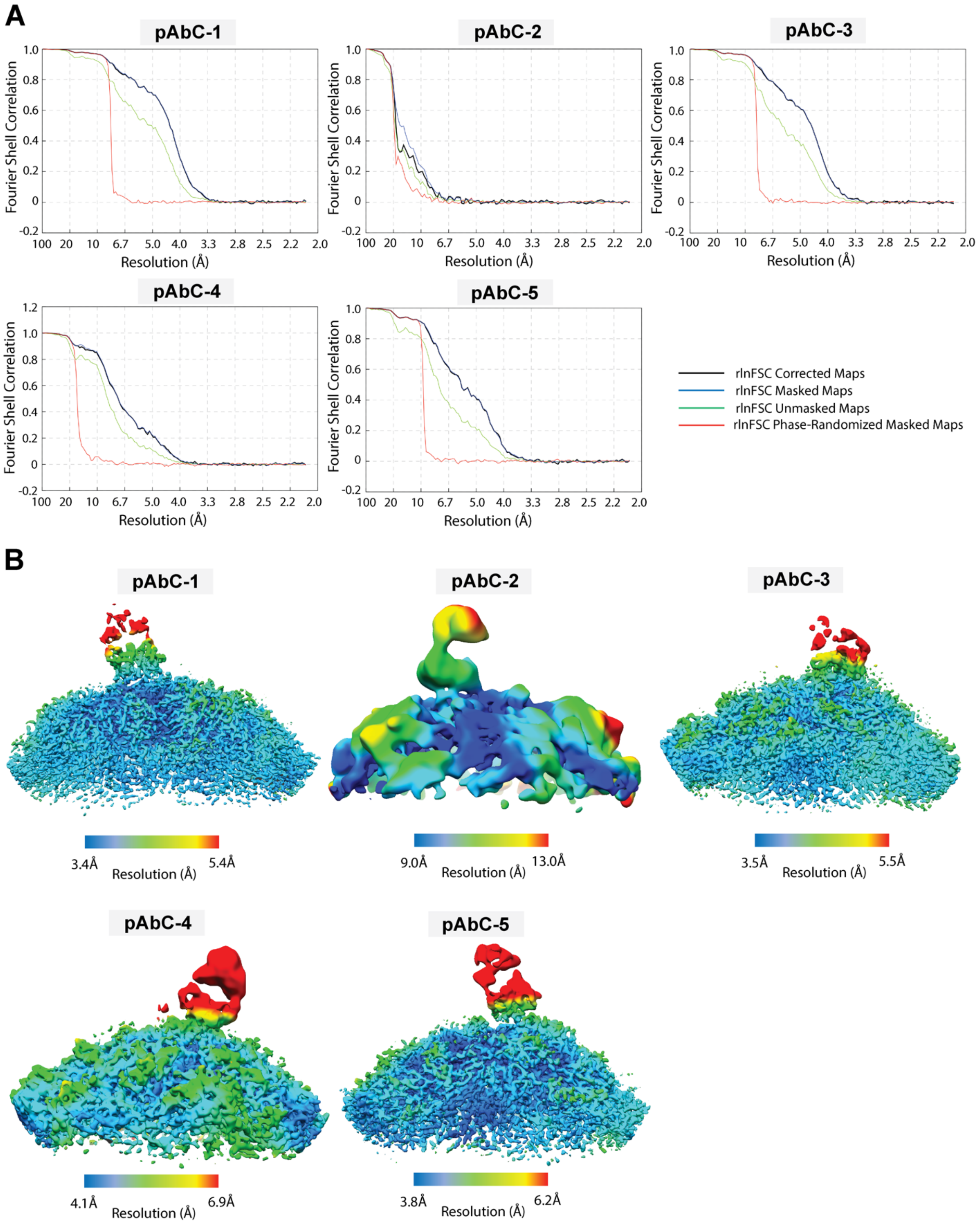


**Figure 2 – figure supplement 4. Extended cryoEMPEM data.** FSC resolution plots **[A]** and local resolution plots **[B]** for EM maps reconstructed by cryoEMPEM analysis of mouse pAb sample. The plots were generated in Relion/3.0.

**Figure 3 – figure supplement 1. Map and model refinement information**

| Complex ID | pAbC-1 | pAbC-2 | pAbC-3 | pAbC-4 | pAbC-5 |
|---|---|---|---|---|---|
| Antigen | CV-A21 | | | | |
| Number of picked particles | 22,937 | | | | |
| Particles after 2D classification | 22,245 | | | | |
| Particles after symmetry expansion | 1,334,700 | | | | |
| Particles in the final map | 19,748 | 7,792 | 29,282 | 22,044 | 12,654 |
| Map symmetry | C1 | C1 | C1 | C1 | C1 |
| Map sharpening B-factor | -65.0 | -100.0 | -55.0 | -40.0 | -45.0 |
| Map Resolution | 3.8 | 12.0 | 3.9 | 4.6 | 4.1 |
| EMDB ID | 26072 | 26069 | 26068 | 26070 | 26071 |
| Residues | | | | | |
| Amino acids | 2758 | N/A | 4451 | N/A | 4418 |
| Ligands (MYR) | 3 | N/A | 5 | N/A | 5 |
| RMSD Bonds (4σ) | 0.020 | N/A | 0.021 | N/A | 0.020 |
| RMSD Angles (4σ) | 1.740 | N/A | 1.743 | N/A | 1.766 |
| Ramachandran | | | | | |
| Outliers (%) | 0.00 | N/A | 0.00 | N/A | 0.00 |
| Allowed (%) | 2.20 | N/A | 1.84 | N/A | 2.22 |
| Favored (%) | 97.80 | N/A | 98.16 | N/A | 97.78 |
| Rotamer outliers (%) | 0.00 | N/A | 0.00 | N/A | 0.00 |
| Clash score | 0.85 | N/A | 1.07 | N/A | 1.60 |
| Molprobity score | 0.81 | N/A | 0.81 | N/A | 0.96 |
| EMRinger score | 3.42 | N/A | 3.31 | N/A | 2.35 |
| PDB ID | 7TQU | N/A | 7TQS | N/A | 7TQT |

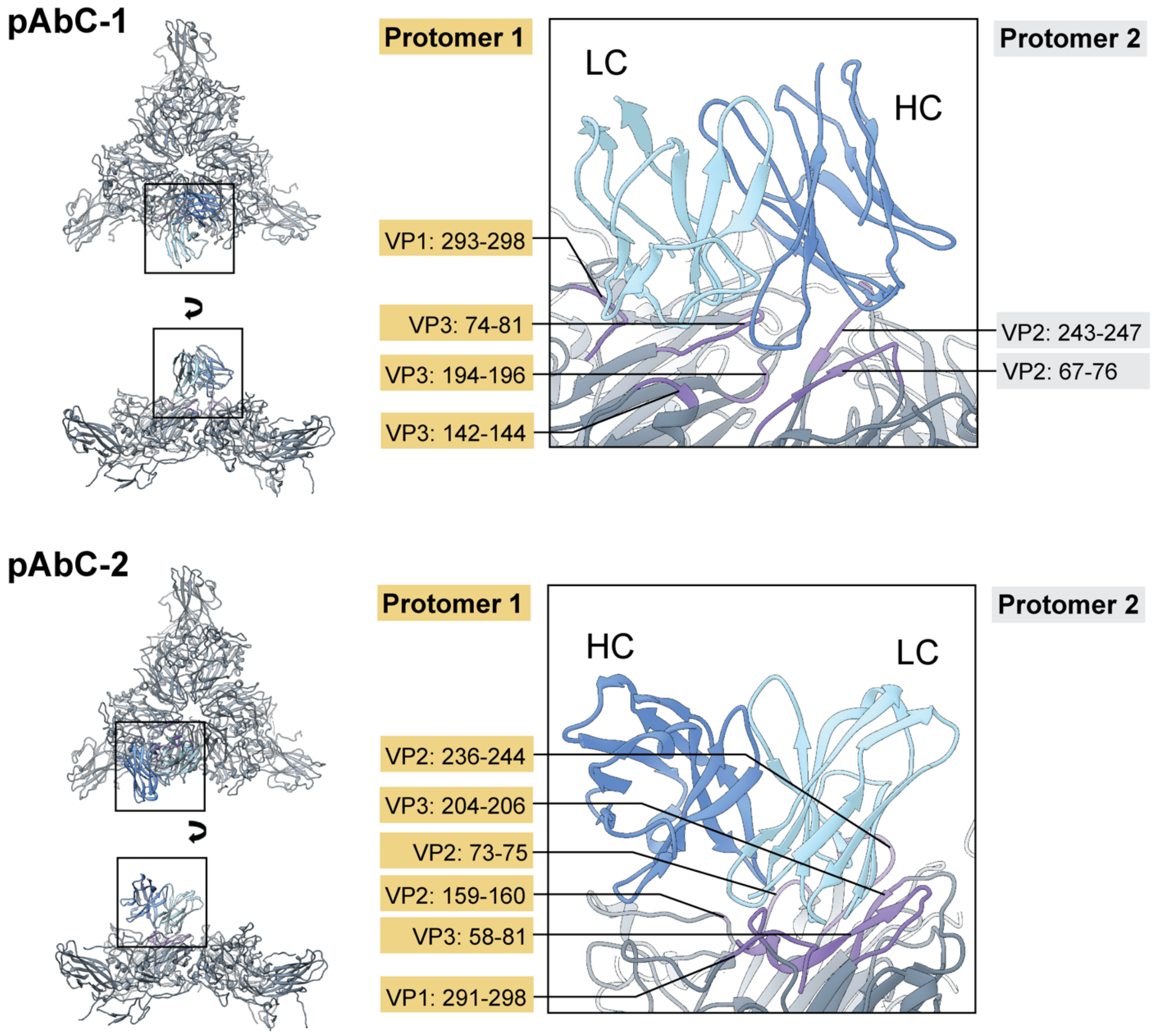


**Figure 3 – figure supplement 2. Epitope-paratope interactions formed by Site-1 targeting polyclonal antibodies, pAbC-1 (top) and pAbC-2 (bottom).** Ribbon representation used throughout the figure. Full models are presented on the left and close-up views of the epitope-paratope interfaces are shown on the right. Heavy and light chains of each antibody are represented in darker and lighter shades of blue, respectively. Contact residues in each epitope are colored purple, while the rest of the antigen is in dark gray. Residue ranges are indicated on the left and right side of the close-up panel and separated based on the protomer they belong to. For pAbC-1 we used a refined model with pAb (Fv fragment) represented as poly-Ala pseudo-model. Map resolution was too low to build a model for pAbC-2 and the presented model was made by docking 3 capsid protomers (each consisting of VP1-4) and a mock mouse Fv fragment (PDB ID: 3i9g) into the pAbC-2 map.

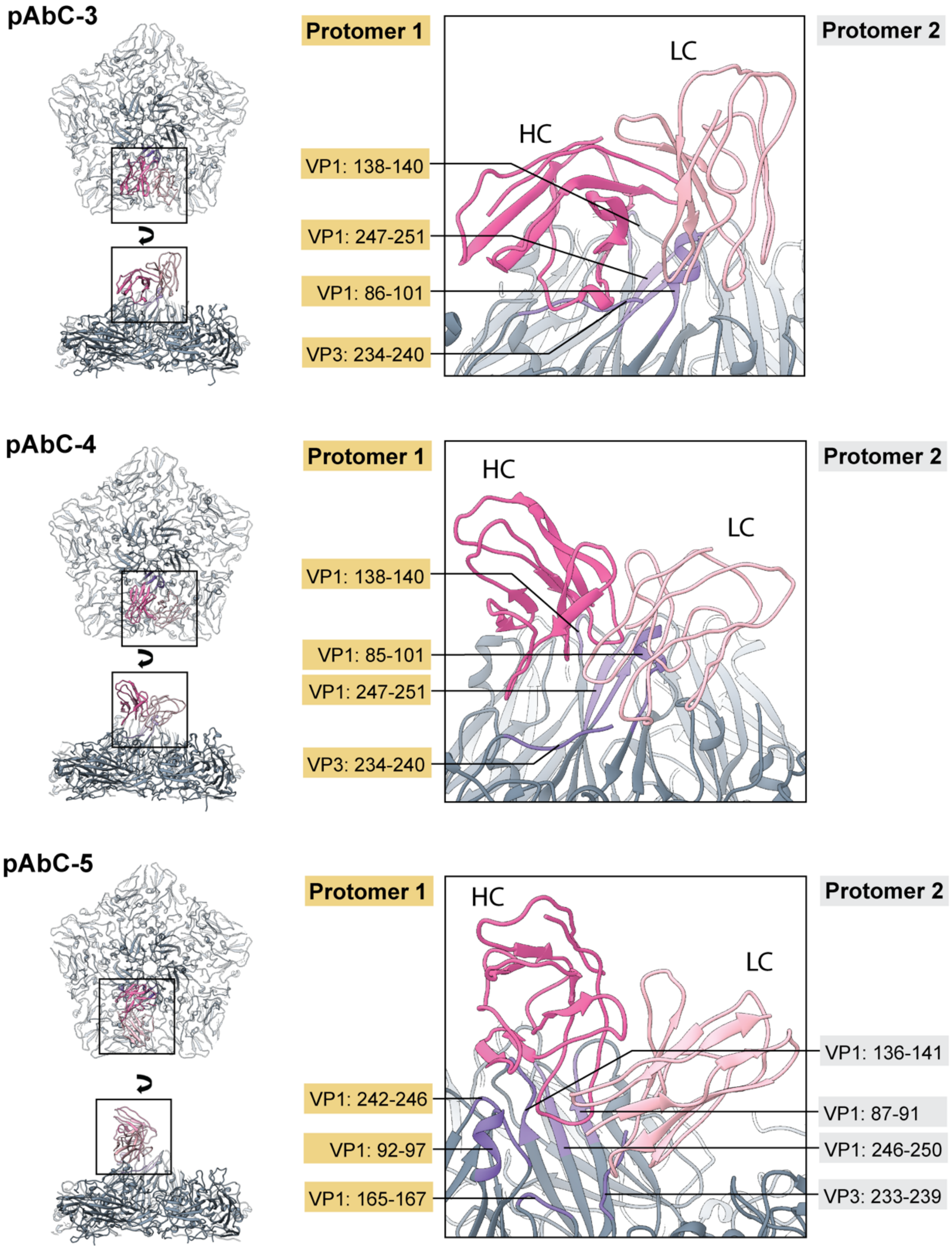


**Figure 3 – figure supplement 3. Epitope-paratope interactions formed by Site-1 targeting polyclonal antibodies, pAbC-3 (top), pAbC-4 (middle) and pAbC-5 (bottom).** Ribbon representation used throughout the figure. Full models are presented on the left and close-up views of the epitope-paratope interfaces are shown on the right. Heavy and light chains of each antibody are represented in darker and lighter shades of pink, respectively. Contact residues in each epitope are colored purple, while the rest of the antigen is in dark gray. Residue ranges are indicated on the left and right side of the close-up panel and separated based on the protomer they belong to. For pAbC-3 and pAbC-5 we used refined models with polyclonal pAbs (Fv fragment) represented as poly-Ala pseudo-models. Map resolution was too low to build a model for pAbC-4 and the presented model was made by docking 5 capsid protomers (each consisting of VP1-4) and a mock mouse Fv fragment (PDB ID: 3i9g) into the pAbC-4 map.

**Figure 3 – figure supplement 4. Epitope contacts within Site-1 and Site-2 (presented as residue ranges)**

| | VP1 | VP2 | VP3 | VP4 |
|---|---|---|---|---|
| **Site-1** | 291-298 | 67-76<br>159-160<br>236-247 | 58-81<br>142-144<br>194-196<br>204-206 | - |
| **Site-2** | 85-101<br>136-141<br>165-167<br>242-251 | - | 233-240 | - |

**Figure 3 – figure supplement 5.** Sequence alignment of capsid-forming polyprotein across enterovirus C CV-A viral strains

```
CV-A21                     MGAQVSTQKTGAHENQNVAANGSTINYTTINYYKDSASNSATRQDLSQDPSKFTEPVKDL  60
CV-A11_(AAQ02676.1)        MGAQVSSQKVGAHENTNVATGGSTVNYTTINYYKDSASNAASKQDFSQDPSKFTEPVKDI  60
CV-A13_(AAQ02677.1)        MGAQVSSQKVGAHENTNVATGGSTVNYTTINYYKDSASNAASKQDFSQDPSKFTEPVKDV  60
CV-A17_(AAQ02679.1)        MGAQVSSQKVGAHENTNVATGGSTVNYTTINYYKDSASNAASKQDFSQDPSKFTEPVKDI  60
CV-A18_(AAQ02680.1)        MGAQVSSQKVGAHENTNVATGGSTVNYTTINYYKDSASNAASKQDFSQDPSKFTEPVKDV  60
CV-A20_(AAQ02682.1)        MGAQVSSQKVGAHENTNVATGGSTVNYTTINYYKDSASNAASKQDFSQDPSKFTEPVKDI  60
CV-A24_(ABM54551.1)        MGAQVSSQKVGAHENTNVATGGSTVNYTTINYYKDSASNAASKQDFSQDPSKFTEPVKDI  60
CV-A1_(AAQ02675.1)         MGAQVSTQKSGSHENQNVAAGGSTINYTTINYYKDSASNSASKQDFSQDPSKFTEPVKDI  60
CV-A19_(AAQ02681.1)        MGAQVSTQKSGSHENQNIAAGGSTINYTTINYYKDSASNSAAKQDFSQDPSKFTEPVKDI  60
CV-A22_(AAQ02683.1)        MGAQVSTQKSGSHENQNIAASGSTINYTTINYYKDSASNSAAKQDFSQDPSKFTEPVKDI  60
                           |-----------------------------VP4--------------------------

CV-A21                     MLKTAPALNSPNVEACGYSDRVRQITLGNSTITTQEAANAIVAYGEWPTYINDSEANPVD  120
CV-A11_(AAQ02676.1)        MLKSAPALNSPNIEACGYSDRVMQLTLGNSTITTQEAANSVVAYGEWPSYLSDKEANPVD  120
CV-A13_(AAQ02677.1)        LIKSAPALNSPNIEACGYSDRVMQLTLGNSTITTQEAANSVVAYGVWPSYLSDKDANPVD  120
CV-A17_(AAQ02679.1)        MLKSAPALNSPNIEACGYSDRVMQLTLGNSTITTQEAANSVVAYGRWPSYLSDREANPVD  120
CV-A18_(AAQ02680.1)        LIKSAPALNSPNIEACGYSDRVMQLTLGNSTITTQEAANSVVAYGVWPSYLSDKDANPVD  120
CV-A20_(AAQ02682.1)        MLKSAPALNSPNVEACGYSDRVLQLTLGNSTITTQEAANSVVGYGQWPTYLNAKDANPVD  120
CV-A24_(ABM54551.1)        MLKSAPALNSPNVEACGYSDRVRQITLGNSTITTQEAANAVVAYGEWPSYLDDREANPVD  120
CV-A1_(AAQ02675.1)         MIKTAPALNSPNIEACGYSDRVLQLTLGNSTITTQEAANSVIAYGEWPSFLSDKEANPVD  120
CV-A19_(AAQ02681.1)        IVKTAPALNSPNIEACGYSDRVLQLTLGNSTITTQEAANAVVAYGEWPRFINDQEANPVD  120
CV-A22_(AAQ02683.1)        MVKTAPALNSPNIEACGYSDRVLQLTLGNSTITTQEAANAVVAYGEWPSFLSDKEANPVD  120
                           ---VP4--|----------------------VP2------------------------

CV-A21                     APTEPDVSSNRFYTLESVSWKTTSRGWWWKLPDCLKDMGMFGQNMYYHYLGRSGYTIHVQ  180
CV-A11_(AAQ02676.1)        QPTEPEVSACRFYTLDTVTWSKSSKGWWWKLPDALKDMGLFGQNMYYHYLGRSGYTVHVQ  180
CV-A13_(AAQ02677.1)        QPTEPDVSACRFYTLDTVEWDKESKGWWWKLPDALKDMGLFGQNMYYHYLGRSGYTVHVQ  180
CV-A17_(AAQ02679.1)        QPTEPDVAACRFYTLESVMWSKESRGWWWKLPDALKDMGLFGQNMFYHYLGRSGYTIHVQ  180
CV-A18_(AAQ02680.1)        QPTEPDVSACRFYTLETVEWDRESKGWWWKLPDALKDMGLFGQNMYYHYLGRSGYTVHVQ  180
CV-A20_(AAQ02682.1)        QPTEPDVSACRFYTLQSVEWKTESKGWWWKLPDALKDMGLFGQNMYYHYLGRSGYTVHVQ  180
CV-A24_(ABM54551.1)        APTEPDVSSNRFYTLDSVQWKSTSRGWWWKLPDALKDMGMFGQNMYYHYLGRSGYTVHVQ  180
CV-A1_(AAQ02675.1)         APTEPDASSNRFYTLDSKPWAADSRGWWWKLPDALKDMGMFGQNMFYHYLGRAGYTVHVQ  180
CV-A19_(AAQ02681.1)        APTEPDASANRFYTLDSVDWGADSKGWWWKLPDALKDMGMFGQNMYYHYLGRAGYTVHVQ  180
CV-A22_(AAQ02683.1)        APTEPDASANRFYTLESITWEKSSRGWWWKLPDALKDMGMFGQNMFYHYLGRAGYTVHVQ  180
                           ------------------------------VP2--------------------------

CV-A21                     CNASKFHQGALGVFLIPEFVMACNTESKTSYVSYINANPGERGGEFTNTYNPSNTDASEG  240
CV-A11_(AAQ02676.1)        CNASKFHQGALGVFAIPEYCMACNTDAKTNYVSYVQANPGEAGGVFTDMYNPS-SETTGA  239
CV-A13_(AAQ02677.1)        CNASKFHQGTLGVFAVPEYCLAGDSNSKTNYTSYVNANPGEAGGKFVSTFTPD-TGTSPK  239
CV-A17_(AAQ02679.1)        CNASKFHQGTLGVFAIPEYCLAGDSDVKNSYTLYVNANPGERGGTFTDKFTAS-SRTNPT  239
CV-A18_(AAQ02680.1)        CNASKFHQGTLGVFAVPEYCLAGDSNSKTNYTSYINANPGERGGTFVSTFTPD-SGAVPK  239
CV-A20_(AAQ02682.1)        CNASKFHQGALGVFAVPEYCLAGDSDVKNSYTTYKNANPGETGGVFVDSFTAT-TQS--T  237
CV-A24_(ABM54551.1)        CNASKFHQGTLGVFAIPEYVMACNTETKTSYVSYVNANPGERGGVFTSTYNPS-TDAAEG  239
CV-A1_(AAQ02675.1)         CNASKFHQGTLLVAAIPEFMMGSNTDTNTGGITYEKANPGEVGGTFQKTATLT-T-GDGK  238
CV-A19_(AAQ02681.1)        CNASKFHQGTLFVAAIPEYMMASNSGTNTGGIIYEFANPGEAGGRFSSTFTPD-TQAPGK  239
CV-A22_(AAQ02683.1)        CNASKFHQGALIVAAIPEFMMGSNTATSTGGVTYANANPGESGGKFSSQFIPS-SDTEAK  239
                           ------------------------------VP2--------------------------

CV-A21                     RKFAALDYLLGSGVLAGNAFVYPHQIINLRTNNSATIVVPYVNSLVIDCMAKHNNWGIVI  300
CV-A11_(AAQ02676.1)        RKFAAVDYLLGCGVLAGNAFVFPHQIINLRTNNCATLVLPYVNSMAIDCMAKHNNWGIAI  299
CV-A13_(AAQ02677.1)        REFQPVDYLFGCGVMAGNAFVFPHQIINLRTNNCATLVLPYVNSLAIDCMAKHNNWGIVI  299
CV-A17_(AAQ02679.1)        RKFCAVDYLLGCGVLAGNAFVFPHQIINLRTNNCATLVLPYVNSLAIDSMTKHNNWGIAI  299
CV-A18_(AAQ02680.1)        REFQPVDYLFGCGVMAGNAFVFPHQIINLRTNNCATLVLPYVNSLAIDCMAKHNNWGIVI  299
CV-A20_(AAQ02682.1)        RKFCPIDYLFGCGVLTGNAFVFPHQIINLRTNNSATLVLPYVNSLAIDCMAKHNNWGLAI  297
CV-A24_(ABM54551.1)        RKFAALDYLLGCGVLAGNAFVFPHQIINLRTNNSATLVLPYVNSLAIDCMAKHNNWGLVI  299
CV-A1_(AAQ02675.1)         NSFCPLDWVLGCGVMAGNATVFPHQFINLRTNNSATLVLPYVNSIATDCMAKHNNWGLVV  298
CV-A19_(AAQ02681.1)        NKFAPLDWLLGCGVMAGNITVFPHQIINLRTNNCATLVLPYVNSVVTDSMAKHNNWGIVV  299
CV-A22_(AAQ02683.1)        NKFAPWDWLLGCGVMAGNITVYPHQIINLRTNNCATLVLPYVNSVATDCMAKHNNWGLVI  299
                           ------------------------------VP2--------------------------
```

**Figure 3 – figure supplement 6.** Sequence alignment of capsid-forming polyprotein across enterovirus C CV-A viral strains (continued)

```
CV-A21                    LPLAPLAFAATSSPQVPITVTIAPMCTEFNGLRNITVPVHQGLPTMNTPGSNQFLTSDDF  360
CV-A11_(AAQ02676.1)       LPLAELDFAEASSPEIPITITIAPMCCEFNGLRNLTSPAKQGLPVMNVPGSNQFLSSDNF  359
CV-A13_(AAQ02677.1)       LPLSKLDYNPDASTKLPITVTIAPMCCEFNGLRNLTIPATQGLPVMSTPGSNQYLTSDNF  359
CV-A17_(AAQ02679.1)       IPLSKLDFAPDASVELPITVTIAPMCCEFNGLRNITIPATQGLPVMNTPGSNQYLTADNF  359
CV-A18_(AAQ02680.1)       LPLSKLDYNPDASTKLPITVTIAPMCCEFNGLRNLTIPATQGLPVMNTPGSNQYLTSDNF  359
CV-A20_(AAQ02682.1)       IPLSKLQFPDTSSTEIPITVTIAPMCCEFNGLRNITVPSTQGLPVMNTPGSNQYLTSDNF  357
CV-A24_(ABM54551.1)       LPLCKLDYAPNSSTEIPITVTIAPMCTEFNGLRNITVPATQGLPTMLTPGSSQFLTSDDF  359
CV-A1_(AAQ02675.1)        MPVVPLQYSNGASTLVPITITIAPMCCEFNGLRSLTTPYTQGLPVMNTPGSNQFLTTDNF  358
CV-A19_(AAQ02681.1)       IPFVKLAYQNGATTKVPITVTIAPMCCEFNGLRSLTAPVTQGLPTMATPGSNQFLTSDNF  359
CV-A22_(AAQ02683.1)       MPFVQLDAVKDATQSVPITVTIAPMCCEFNGLRSLTVPVLQGLPTMSTPGSNQFLTSDDF  359
                          --------------------VP2----------------|---------VP3-------

CV-A21                    QSPCALPNFDVTPPIHIPGEVKNMMELAEIDTLIPMNAVDGKVNTMEMYQIPLNDNLSKA  420
CV-A11_(AAQ02676.1)       QSPCALPEFDVTPPIHIPGEVRNMMELAEIDTLIPMDLSESKKNTMGMYRVELGSG-KSL  418
CV-A13_(AAQ02677.1)       QSPCALPEFDVTQPIFIPGEVKNLMELAEIDTMIPMDLSEGKRNSMDMYRVKISDA-GDR  418
CV-A17_(AAQ02679.1)       QSPCALPEFDVTQPIDIPGEVKNLMEIAEIDTMIPLDLSESKKNTMDMYRVQLQASPSNR  419
CV-A18_(AAQ02680.1)       QSPCALPEFDVTQPIFIPGEVKNLMELAEIDTMIPMDLSEGKKNTMEMYRVKLSDT-GNR  418
CV-A20_(AAQ02682.1)       QSPCALPEFDVTQAINIPGEVKNIMEIAEIDTMIPLNLSDSRKNSMDMYRVPVTTS-ADL  416
CV-A24_(ABM54551.1)       QSPCALPNFDVTPPIHIPGEVTNMMELAEIDSMIPMNSVTGKANTMEMYPIPLNDK-GSA  418
CV-A1_(AAQ02675.1)        QSPCALPDFDVTPEIHIPGEVKNMMELAEIDSLVPMNAVAGKVNSMEAYQIPIQANQQDN  418
CV-A19_(AAQ02681.1)       QSPCALPDFDVTPEIHIPGEVKNMMELAEIDTLIPMNAIAKKVDTMEAYPIPLQAGVQNN  419
CV-A22_(AAQ02683.1)       QSPCALPNFDVTPAIHIPGEVKNMMELAEIDTLIPMNAISTNVNKMEAYPIPLQASKPNA  419
                          ------------------------------VP3---------------------------

CV-A21                    --PIFCLSLSPASDKRLSHTMLGEILNYYTHWTGSIRFTFLFCGSMMATGKLLLSYSPPG  478
CV-A11_(AAQ02676.1)       SKPILCLSLSPASEQRLGYTMLGEILNYYTHWSGSLKFTFLFCGSMMATGKILISYAPPG  478
CV-A13_(AAQ02677.1)       NKPILCLTLSPASDPRLSYTMLGEILNYYTHWAGSLKFSFLFCGSMMATGKLLVAYSPPG  478
CV-A17_(AAQ02679.1)       DTPILCLSLSPASDPRLSFTMLGEILNYYTHWAGSLKFSFLFCGSMMATGKILVSYAPPG  479
CV-A18_(AAQ02680.1)       DKPILCLSLSPASDPRLSYTMLGEILNYYTHWAGSLKFSFLFCGSMMATGKLLVAYSPPG  478
CV-A20_(AAQ02682.1)       DKPILCLSLSPASDERLSYTMLGEILNYYTHWAGSIKYTFLFCGSMMATGKLLIAYAPPG  476
CV-A24_(ABM54551.1)       D-PIFSISLSPASDKRLQYTMLGEILNYYTHWTGSLRFTFLFCGSMMATGKILLSYSPPG  477
CV-A1_(AAQ02675.1)        NKSIFAISLSPAANPRLSYTMLGEILNYYTHWSGSIKFTFLFCGSAMATGKLLLSYSPPG  478
CV-A19_(AAQ02681.1)       NQSIFSISLSPAADQRLSRTMLGEILNYYTHWTGSIKFTFLFCGSMMATGKILLSYSPPG  479
CV-A22_(AAQ02683.1)       SESIFSISLSPAADERLAHTMLGEILNYYTHWTGSLKFTFLFCGSMMATGKILISYSPPG  479
                          ------------------------------VP3---------------------------

CV-A21                    AKPPTNRKDAMLGTHIIWDLGLQSSCSMVAPWISNTVYRRCARDDFTEGGFITCFYQTRI  538
CV-A11_(AAQ02676.1)       AKPPTTRREAMLGTHVIWDIGLQSSATMVVPWISNVMYRRCVKDDFTEGGYISMFYQTKI  538
CV-A13_(AAQ02677.1)       AQPPQDRKAAMLGTHVIWDIGLQSSCTMVVPWISNTSYRRTVKDDFTEGGYISMFYQTRV  538
CV-A17_(AAQ02679.1)       AQPPKTRKDAMLGTHLIWDIGLQSSCTMVVPWISNTAYRRTIKDDFTEGGYISMFYQTKI  539
CV-A18_(AAQ02680.1)       AQPPQDRKAAMLGTHVIWDIGLQSSCTMVVPWISNTSYRRTAKDDFTEGGYISMFYQTRI  538
CV-A20_(AAQ02682.1)       AKPPRTRKEAMLGTHVIWDVGLQSSCTMVVPWISNTAYRRTVEDDFTEGGYISMFYQTKI  536
CV-A24_(ABM54551.1)       ASPPKTRKDAMLGTHIIWDLGLQSSCTMLAPWISNTVYRRCVKDDFTEAGYITCFYQTRI  537
CV-A1_(AAQ02675.1)        AKPPTTRKEAMLGTHIIWDVGLQSSATMVAPWISNVNYRRCVKDDFTEGGYICCFYQTAI  538
CV-A19_(AAQ02681.1)       AKPPTTRKEAMLGTHLIWDIGLQSSATMVAPWISNVNYRRCVKDDFTEGGYICCFYQTAI  539
CV-A22_(AAQ02683.1)       AKPPTTRKEAMLGTHVIWDLGLQSSVTLVAPWISNVNYRRCVRDDFTEGGYICAFYQTAI  539
                          ------------------------------VP3---------------------------

CV-A21                    VVPASTPTSMFMLGFVSACPDFSVRLLRDTPHISQSKLIGRTQG-IEDLIDTAIKNALRV  597
CV-A11_(AAQ02676.1)       VVPLSTPTTMTLLSFVSACNDFTVRLLRDTTHISQTTK-INTQGPIEEIISTVASNALAL  597
CV-A13_(AAQ02677.1)       VVPASTPTSMDILCFISACNDFTVRLLRDTTHITQSAL---PQG-LEDLIQQVASNALQL  594
CV-A17_(AAQ02679.1)       VVPASTPTTMDIIGFVSACNDFSVRLLRDTTHIAQTAM---PQG-IEDLIQQVASNALQI  595
CV-A18_(AAQ02680.1)       VVPASTPTSMDILCFVSACNDFTVRLLRDTTHISQSAM---PQG-LEDLIQQVATNALSL  594
CV-A20_(AAQ02682.1)       VVPASTPTNMDILGFVSACNDFSVRLLRDTTHISQTAM---PQG-IEDLITEVASNALKL  592
CV-A24_(ABM54551.1)       VVPSSTPTSMFMLAFVSACPDFSVRLLQDTSHISQTALVARTQG-IEDTIDTVINNALQL  596
CV-A1_(AAQ02675.1)        VVPSGTPTVTMSMLCFVSACNDFTARLLKDSPHVIQNNAV--AQG-LGDSIEAAIDSITQN  595
CV-A19_(AAQ02681.1)       VVPPGAPTDMSMLAFVSACNDFSARLLKDTPFITQQALTVTTQG-IDDIIDNVVTNALKV  598
CV-A22_(AAQ02683.1)       IVPPSTPTAMYMLAFVSACNDFSVRLLKDTPFVRQDQY-VNTQG-IEDTIEKVVGDALRV  597
                          --------------------VP3------------------|-------VP1-------
```

**Figure 3 – figure supplement 7.** Sequence alignment of capsid-forming polyprotein across enterovirus C CV-A viral strains (continued)

```
CV-A21                    SQP---------PSTQSTEATSGVNSQEVPALTAVETGASGQAIPSDVVETRHVVNYKTR  648
CV-A11_(AAQ02676.1)       SQP-KPV----DNSVQNTQQSAPVHSQEVPALTAVETGATSDVVPSDLIQTRHVLNVKSR  652
CV-A13_(AAQ02677.1)       SQPTRPALPPAEQSVPNTNQTTPEHSKEVPALTAVETGATNPLEPGDTVQTRHVIQTRSR  654
CV-A17_(AAQ02679.1)       SQPTRPALPSTE-SLPNTQQSAPSHSQEVPALTAVETGATNPLEPSDTVQTRHVIQTRSR  654
CV-A18_(AAQ02680.1)       SQPTRPALPPAEQSVPNTSQTTPEHSKEVPALTAVETGATNPLEPGDTVQTRHVVQTRSR  654
CV-A20_(AAQ02682.1)       SQP-KPS---TQQSLPNTSSSEPTHSQEAPALTAVETGATSSVVPADLVQTRHVIQTRSR  648
CV-A24_(ABM54551.1)       SQP----QPNKQLTAQSTPSTSGVNSQEVPALTAVETGASGQAVPSDVIETRHVVNYKTR  652
CV-A1_(AAQ02675.1)        ALT----------TVQNTTQSGPTHSKEVPALTAVETGATSQVEPGDLIETRHVINMRQR  645
CV-A19_(AAQ02681.1)       SMP----------QVQDTQSSGPVNSKEVPALTAVETGATSQVDPSDLIETRHVINNRLR  648
CV-A22_(AAQ02683.1)       SMP----------QVANTQPSGPVNSKEVPALTAVETGATSQVTPEDLIETRHVINNRLR  647
                          ------------------------------VP1---------------------------

CV-A21                    SESCLESFFGRAACVTILSLTNSSKS---GEEKKHFNIWNITYTDTVQLRRKLEFFTYSR  705
CV-A11_(AAQ02676.1)       SESTIESFFARAACVTIMQVDNFNAT-SVEDKRKLFAKWAITYTDTVQLRRKLEFFTYSR  711
CV-A13_(AAQ02677.1)       SESTVESFFARGACVTIMGVDNYNETLKGDQKSTLFTTWNITYTDTVQLRRKLEMFTYSR  714
CV-A17_(AAQ02679.1)       SESTIESFFARGACVTIMTVENFNAT-EAADKKKLFATWNITYTDTVQLRRKLEMFTYSR  713
CV-A18_(AAQ02680.1)       SESTVESFFARGACVTIMGVDNYNESLTSSQKSTLFATWNITYTDTVQLRRKLEMFTYSR  714
CV-A20_(AAQ02682.1)       SESTVESFFARGACVTIMSVENYNET--AIAESKLFTKWNITYTDTVQLRRKLEMFTYSR  706
CV-A24_(ABM54551.1)       SESTLESFFGRSACVTIIEVENFNAT-SEADKRKQFTTWPITYTNTVQLRRKLEFFTYSR  711
CV-A1_(AAQ02675.1)        SEASIESFFGRSACVAILGLSNAKPT--DTNTKQLFKTWRISYLETHQLRRKLEFFTYSR  703
CV-A19_(AAQ02681.1)       SECTIESFFGRSACVAIIGLSNQKPT--SDNAAKLFATWKISYLDMYQLRRKLEFFTYSR  706
CV-A22_(AAQ02683.1)       SECTVEAFFGRSACVAILGVVNKKPD--TTNAKDLFTTWRITYLQTYQLRRKLELFTYSR  705
                          ------------------------------VP1---------------------------

CV-A21                    FDLEMTFVFTENYPSTASGEVRNQVYQIMYIPPGAPRPSSWDDYTWQSSSNPSIFYMYGN  765
CV-A11_(AAQ02676.1)       FDLEMTFVLTERYYSQSSGHARSQVYQIMYVPPGAPTPSAWDDYTWQTSSNPSIFFTTGN  771
CV-A13_(AAQ02677.1)       FDIEFTFVVTERYYSSNSGHALNQVYQIMYVPPGAPVPKKWDDYTWQTSSNPSIFYTYGS  774
CV-A17_(AAQ02679.1)       FDIEFTFVTTERYYASNSGHARNQVYQLMYVPPGAPVPQQWDDYTWQTSSNPSVFYTYGD  773
CV-A18_(AAQ02680.1)       FDIEFTFVVTERYYSSNSGHALNQVYQIMYVPPGAPIPKKWDDYTWQTSSNPSIFYTYGT  774
CV-A20_(AAQ02682.1)       FDIEFTFVVTERYHSANSGHALNQVYQIMYVPPGAPVPQRWDDYTWQTSSNPSVFYTYGT  766
CV-A24_(ABM54551.1)       FDLEMTFVVTERYYASNTGHARNQVYQIMYIPPGAPQPTAWDDYTWQSSSNPSVFYTYGS  771
CV-A1_(AAQ02675.1)        FDLEMTIVITERVFNAVNVPLRNYVYQIMYVPPGAPEPQSWDDYTWQSSTNPSIFYTTGN  763
CV-A19_(AAQ02681.1)       FDLELTFVISERFFTSTSAAARDYVYQIMYIPPGAPIPQVWDDYTWQSSTNPSIFYTTGN  766
CV-A22_(AAQ02683.1)       FDLELTFVITERYFSGTAATTRDYVYQIMYVPPGAPIPNTWDDYTWQSSTNPSVFYTTGN  765
                          ------------------------------VP1---------------------------

CV-A21                    APPRMSIPYVGIANAYSHFYDGFARVPLEGENTDAGDTF-YGLVSINDFGVLAVRAVNRS  824
CV-A11_(AAQ02676.1)       APPRISIPFVGIANAYSHFYDGFSRVPLEGETTDTGDAY-YGLTSINDFGTLAVRVVNDY  830
CV-A13_(AAQ02677.1)       APPRISIPFVGIANAYSHFYDGYATVPLKTDTTDSGAAY-YGAVSINDFGLLAVRVVNEH  833
CV-A17_(AAQ02679.1)       APARISIPFVGIANAYSHFYDGYAVVPLKDSTQDAGAAY-YGATSINDFGMLAVRVVNEF  832
CV-A18_(AAQ02680.1)       APPRISIPFVGITNAYSHFYDGYATVPLKTDTTDPGAAF-YGAVSINDFGLLAVRVVNEH  833
CV-A20_(AAQ02682.1)       APARISIPYVGIANAYSHFYDGFAKVPIEGETSDPGDAY-YGATSINDFGILAIRVVNEH  825
CV-A24_(ABM54551.1)       APPRMSIPYVGIANAYSLFYDGFARVPLKDETADSGDTF-YGLVTINDFGILAIRVVNEF  830
CV-A1_(AAQ02675.1)        APPRVSIPFVGIGSAYSHFYDGFSQIPL--DSISAGASNKYGYTSINDFGTLAIRIVNEY  821
CV-A19_(AAQ02681.1)       ACPRVSIPFVGIGAAYSHFYDGFSLVPF--NTIDAGASNRYGYTTINDFGTMAIRIVNEY  824
CV-A22_(AAQ02683.1)       ASPRMSIPFVGIGAAYAHFYDGFSVVPF--NQIDAGASNKYGYSSIKDFGTLAVRIVNEF  823
                          ------------------------------VP1---------------------------

CV-A21                    NPHTIHTSVRVYMKPKHIRCWCPRPPRAVLYRGEGVDMISSAILPLAKVDSITTF
CV-A11_(AAQ02676.1)       NPARVETRIRVYMKPKHVRVWCPRPPRAVSYRGPGVDLLSTSVTPLSK-HDLATY
CV-A13_(AAQ02677.1)       NPVRVSSKIRVYMKPKHVRVWCPRPPRAVEYYGPGVDYKANTLTPLPI-KNLTTY
CV-A17_(AAQ02679.1)       NPARITSKLRVYMKPKHVRVWCPRPPRVVPYFGPGVDYK-DSLTPLST-KALNTY
CV-A18_(AAQ02680.1)       NPVRVSSKIRVYMKPKHVRVWCPRPPRAVEYYGPGVDYKANTLTPLPT-KNLTTY
CV-A20_(AAQ02682.1)       NPVQVSSKIRVYMKPKHVRVWCPRPPRAVPYFGPGVDYKGDALTPLSR-KDLTTY
CV-A24_(ABM54551.1)       NPARITSKIRVYMKPKHVRCWCPRPPRAVPYRGEGVDFNSSSITPLTAVANINTF
CV-A1_(AAQ02675.1)        DPVQVDAKARVYIKPKHVRMWCPRPPRAMPYKNSTVDFDPSATV-MTQVADIRTY
CV-A19_(AAQ02681.1)       DPVTIDAKVRVYMKPKHIKVWCPRPPRAVAYNGPTVNFNENPHV-MTAVADIRTY
CV-A22_(AAQ02683.1)       DPVTIEAKVRVYMKPKHVRVWCPRPPRAVPYQNSSVDFAQNAVA-MNQVATIRTY
                          ---------------------------VP1-------------------------
```

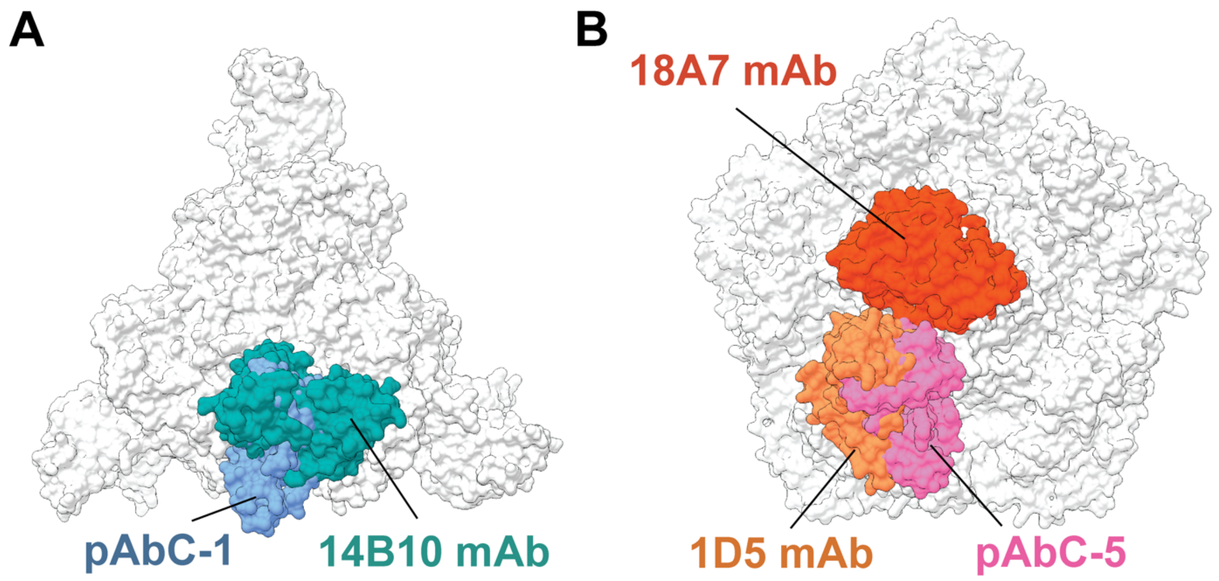


**Figure 3 – figure supplement 8. Structural comparison of published CV-specific monoclonal antibodies and polyclonal antibodies recovered in this study. [A]** Overlay of the structures of Site-1 targeting antibodies, pAbC-1 and 14B10 mAb (ref). **[B]** Overlay of the structures of Site-2 targeting antibodies, pAbC-5, 1D5 (ref) and 18A7 (ref).
